# Fluctuating light experiments and semi-automated plant phenotyping enabled by self-built growth racks and simple upgrades to the IMAGING-PAM

**DOI:** 10.1101/795476

**Authors:** Dominik Schneider, Laura S. Lopez, Meng Li, Joseph D. Crawford, Helmut Kirchhoff, Hans-Henning Kunz

## Abstract

**Background:** Over the last years, several plant science labs have started to employ fluctuating growth light conditions to simulate natural light regimes more closely. Many plant mutants reveal quantifiable effects under fluctuating light despite being indistinguishable from wild-type plants under standard constant light. Moreover, many subtle plant phenotypes become intensified and thus can be studied in more detail. This observation has caused a paradigm shift within the photosynthesis research community and an increasing number of scientists are interested in using fluctuating light growth conditions. However, high installation costs for commercial controllable LED setups as well as costly phenotyping equipment can make it hard for small academic groups to compete in this emerging field.

**Results:** We show a simple do-it-yourself approach to enable fluctuating light growth experiments. Our results using previously published fluctuating light sensitive mutants, *stn7* and *pgr5*, confirm that our low-cost setup yields similar results as top-prized commercial growth regimes. Moreover, we show how we increased the throughput of our Walz IMAGING-PAM, also found in many other departments around the world. We have designed a Python and R-based open source toolkit that allows for semi-automated sample segmentation and data analysis thereby reducing the processing bottleneck of large experimental datasets. We provide detailed instructions on how to build and functionally test each setup.

**Conclusions:** With material costs well below USD$1000, it is possible to setup a fluctuating light rack including a constant light control shelf for comparison. This allows more scientists to perform experiments closer to natural light conditions and contribute to an emerging research field. A small addition to the IMAGING-PAM hardware not only increases sample throughput but also enables larger-scale plant phenotyping with automated data analysis.

## Background

In nature, plants frequently experience rapidly changing light conditions. This phenomenon is mainly caused by shading effects within the plant canopy or between neighboring plants. Additionally, cloud movements and pollutants cause changes in the light quality and quantity (Slattery et al., 2018). Plants have evolved several molecular mechanisms for coping with light stress of which the most important one is non-photochemical quenching (NPQ) (Jahns and Holzwarth, 2012). NPQ protects the plant effectively during high light by dissipating light energy as heat rather than allowing the energy to be put towards photochemistry. However, plants rapidly deactivate NPQ to maximize productivity when light availability becomes limiting. A number of enzymes and transport proteins critical in this process have been discovered over the last years (Armbruster et al., 2017). This research progress was mainly achieved by switching from constant to dynamic growth lights mimicking natural conditions. More researchers should employ dynamic growth regimes to address open questions, but professional chambers with controllable LED elements and tools to determine photosynthesis come at a high cost.

Pulse-Amplitude-Modulation (PAM) chlorophyll fluorescence measurements represent a centerpiece of fitness evaluation for plants, algae and cyanobacteria (Brooks and Niyogi, 2011). Although primarily aimed at providing quantitative insight into the photosynthetic light reactions, several parameters determined during the measurements were found to be reliable indicators of a plant’s response towards abiotic and biotic stresses (Murchie and Lawson, 2013). Notably, chlorophyll fluorometers are frequently used detectors in automated phenotyping platforms. However, automated phenotyping requires significant investment and therefore platform installations and usage remain limited to few institutions worldwide.

Since its release in the mid-2000s, the IMAGING-PAM, a manual bench-top camera-based chlorophyll fluorometer sold by Walz GmbH, has been widely applied in various types of research on phototropic organisms around the world (Escher et al., 2006). A brief Google scholar inquiry using the search term “IMAGING-PAM” yielded over 2300 results. Even though the machine offers many useful features, sample throughput and downstream data analysis are slow and cumbersome. These limitations have made it difficult to apply the IMAGING-PAM in larger scale experiments which are needed to unveil more subtle performance differences with low statistical power and for screening mutant or germplasm collections under an ever-increasing variety of treatment conditions. Experiment complexity and size are further expanded when previously published mutants are included as reference points.

Downstream data processing can benefit greatly from making subtle hardware adjustments. Consistent sample positioning and image capture settings facilitate scriptable image analysis tools (Tovar et al., 2018). Since no standardized imaging setup exists for the IMAGING-PAM, we addressed the issue by designing an easy-to-build sample holder kit which enables straight-forward plant handling and guarantees consistent and reproducible positioning of individuals between experiments. Together these changes improve picture quality, increase sample throughput, and enable a more automated downstream data analysis pipeline.

## Results

### Order parts to build a low-cost plant growth rack for fluctuating light experiments

Initially, all parts were purchased online. Table 1 summarizes each manufacturer and the item numbers. The items and pricing represent a loose guideline and might be outdated at the time of reading this article. Parts by other manufacturers may work just as well and may provide even cheaper options. However, the parts listed were thoroughly tested in this study and all parts work well together.

**Table 1:**
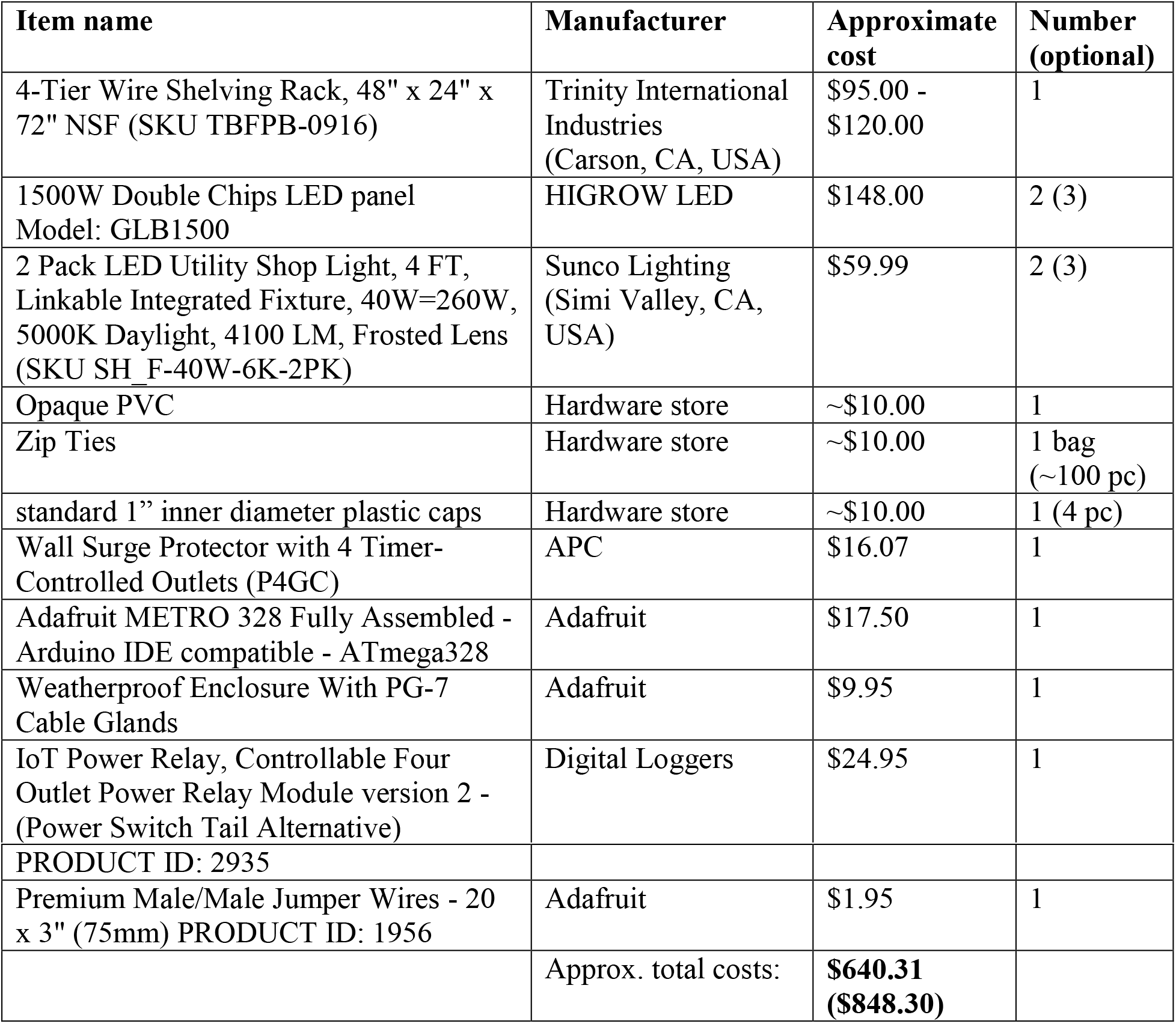
Parts needed for construction of fluctuating light plant growth rack

| Item name | Manufacturer | Approximate cost | Number (optional) |
| --- | --- | --- | --- |
| 4-Tier Wire Shelving Rack, 48" x 24" x 72" NSF (SKU TBFPB-0916) | Trinity International Industries (Carson, CA, USA) | \$95.00 - \$120.00 | 1 |
| 1500W Double Chips LED panel<br>Model: GLB1500 | HIGROW LED | \$148.00 | 2 (3) |
| 2 Pack LED Utility Shop Light, 4 FT, Linkable Integrated Fixture, 40W=260W, 5000K Daylight, 4100 LM, Frosted Lens (SKU SH_F-40W-6K-2PK) | Sunco Lighting (Simi Valley, CA, USA) | \$59.99 | 2 (3) |
| Opaque PVC | Hardware store | ~\$10.00 | 1 |
| Zip Ties | Hardware store | ~\$10.00 | 1 bag (~100 pc) |
| standard 1" inner diameter plastic caps | Hardware store | ~\$10.00 | 1 (4 pc) |
| Wall Surge Protector with 4 Timer-Controlled Outlets (P4GC) | APC | \$16.07 | 1 |
| Adafruit METRO 328 Fully Assembled - Arduino IDE compatible - ATmega328 | Adafruit | \$17.50 | 1 |
| Weatherproof Enclosure With PG-7 Cable Glands | Adafruit | \$9.95 | 1 |
| IoT Power Relay, Controllable Four Outlet Power Relay Module version 2 - (Power Switch Tail Alternative) | Digital Loggers | \$24.95 | 1 |
| PRODUCT ID: 2935 |  |  |  |
| Premium Male/Male Jumper Wires - 20 x 3" (75mm) PRODUCT ID: 1956 | Adafruit | \$1.95 | 1 |
| | Approx. total costs: | **\$640.31**<br>**(\$848.30)** | |

### Setup of a low-cost plant growth rack for dynamic light experiments

Initially, the wire shelving rack was assembled with three levels according to the manufacturer’s instructions. The distance between the shelves’ lowest to highest point was 39 cm (Figure 1A). Hanging from the middle shelf, 2-40W LED grow lights provide constant light and were affixed using zip ties. It is important to use LED grow lights that can be connected in series as this simplifies the control of the entire rack. Additionally, these lights should output a broadband light spectrum similar to the sun. The two light fixtures were hung on the most outside position and had a distance of 29.5 cm to each other (Figure 1B). Light intensities on the Arabidopsis leaf rosette level were found to be consistent around 90 μmol photons m^−2^ s^−1^ with a leaf surface temperature of 23.9°C ± 0.5. The capacity of our constant light setup is 200 2” x 2” x 2⅛” (5 cm x 5 cm x 5.5 cm) pots that are ideal for growing single Arabidopsis plants.

**Figure 1.**
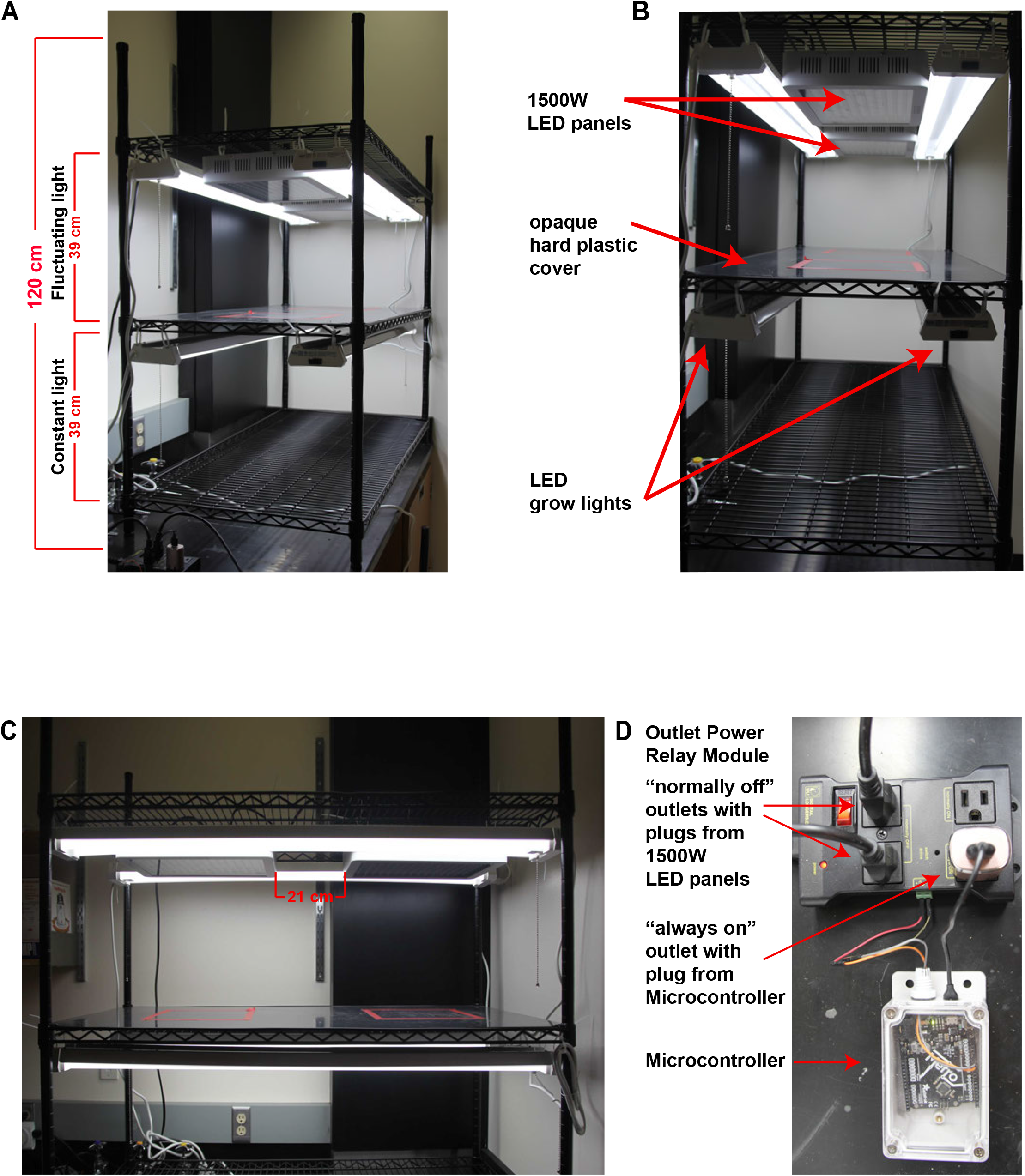
Design of low-cost fluctuating light plant rack. A) Front view of the growth rack (120 cm total height). Constant light section at the bottom and fluctuating light section above with a height of 39 cm each. B) In both sections two daisy-chained LED grow lights were placed 29.5 cm apart from each other. Additionally, in the FL section, two daisy-chained 1500W LED panels were installed 21 cm away from each other. An opaque hard-plastic cover divides the FL from the constant light section. C) Side-view of the rack. D) The 1500W LED panels are plugged into a controllable outlet power relay module controlled by a micro-controller, which determines when the panels turn on and off (1 min at 900 μmol photons m^−2^ s^−1^ and 4 min at 90 μmol photons m^−2^ s^−1^). The outlet power relay module and the LED shop lights run on timer-controlled outlets that keep both units on for 12hrs.

Another pair of LED grow lights was installed similarly one shelf above to function as the background light for a fluctuating light system. Both LED grow light units were individually plugged into a surge-protected power strip with integrated timer function set to 12 h on from 8 AM to 8 PM. Between the upper background lights, two broad spectrum 1500 W LED panels were positioned and strapped onto the rack using zip ties (Figure 1B-C). The distance between the two panels was 21 cm. These two 1500 W LED units were also cable-connected with each other. The single cable from the 1500 W LED panel unit was plugged into one of the “normally off’ outlets in the controllable Outlet Power Relay Module (Figure 1D). Light intensities on the Arabidopsis leaf rosette level are on average 900 μmol photons m^−2^ s^−1^ when both the background LEDs and the two 1500 W LED panels run simultaneously with a leaf surface temperature of 27.3°C ± 1.0 at the end of a one minute high light period. The entire installation should be inspected by a certified electrician to ensure the unit complies with local safety standards. The capacity of our fluctuating light setup is 90 2” x 2” x 2⅛” (5 cm x 5 cm x 5.5 cm) pots. This number is reduced from the lower shelf because the 1500W LED units provide a smaller swath of illumination compared to the LED grow lights. One disadvantage of the low-priced LED panels is that their light intensity cannot be implicitly changed. Changes to the light intensity would require an additional voltage regulator, LED panels with different wattage, or adjusting the distance between the panels and the plants.

A rigid, dark, and opaque hard plastic cover was cut and put on the middle shelf to protect plants on the lower shelf from the high light intensities above. The plastic cover also prevents water spill into the electric equipment below. Lastly, the posts were cut off right above the shelf holding the two 1500 W LED panels. All new ends should be filed down and capped to avoid injuries. Because the 1500 W LED panels produce heat and have fan openings, it is not safe to use the space directly above. This safety precaution also guided our decision to install the fluctuating light system in the upper half of the shelving.

The remaining post pieces (~65 cm length) and the last wire shelf were later used to build a smaller, secondary growth rack by adding an additional set of LED grow lights and one additional 1500 W LED panel with an opaque divider in the middle of the shelf (Additional file 1A). We used the same Outlet Power Relay Module so we were able to increase our capacity (27 additional plants under fluctuating light and 50 additional plants under constant light) for minimal additional cost (Table 1).

A simple Adafruit micro-controller was connected to the Outlet Power Relay Module to control the light pulses (i.e. output from the 1500 W LED panels). It was flashed with a script (Additional file 2) that turns on the “normally off” outlet every 5 minutes for exactly one minute (Figure 1D). Therefore, the plants become exposed to alternating high light (1 min at 900 μmol photons m^−2^ s^−1^) and low light (4 min at 90 μmol photons m^−2^ s^−1^) (Additional file 1B). Minor adjustments to the script could enable other light pulse frequencies or durations. The microcontroller itself receives its power via the “always on” outlet on the Power Relay Module. The Power Relay Module was connected to the timer-controlled power strip (12 h on from 8 AM to 8 PM). To protect the micro-controller unit from moisture it is strongly advisable to use a weatherproof enclosure.

### Testing the fluctuating light plant growth rack using known loss-of-function mutants

Among the best described Arabidopsis mutants susceptible to fluctuating light are *stn7* and the *pgr5* loss-of-function mutants. While *stn7* shows strongly diminished growth under fluctuating light, *pgr5* is even more sensitive to the same conditions and dies rapidly after being shifted into fluctuating light (Tikkanen et al., 2010). Therefore, both loss-of-function lines serve as ideal controls to test how closely the newly constructed growth rack reproduces previously published results from independent international research groups.

STN7 represents a thylakoid serine-threonine protein kinase that phosphorylates Light Harvesting Complex (LHC) II to allow for migration of the complex from photosystem II (PSII) to PSI. The lack of this kinase therefore renders *stn7* loss-of-function mutant unable to adapt to changing light conditions adequately (Bellafiore et al., 2005; Bonardi et al., 2005). First, *stn7* and WT were germinated and grown in 12/12 h day-night cycles using constant lighting (90 μmol photons m^−2^ s^−1^) on the lower shelf. At an age of 14 days, half the plants from each genotype remained on the lowest shelf whereas the other half was moved onto the upper shelf where plants were exposed to the previously described fluctuating light conditions (1 min at 900 μmol photons m^−2^ s^−1^, 4 min at 90 μmol photons m^−2^ s^−1^; 12/12 h day-night cycles at room temperature ~24°C). At a plant age of four weeks, size differences between the two light treatments became clearly visible. There was no growth difference between the genotypes under constant light, but *stn7* revealed visually less leaf surface than WT under fluctuating light (Figure 2A). Both observations are in line with previously reported characteristics of *stn7* (Tikkanen et al., 2010; Grieco et al., 2012). Additionally, when photosynthesis-related parameters of dark-adapted plants were determined, *stn7* revealed reduced *F*_v_/*F*_m_ values (Maximum quantum yield of PSII (Maxwell and Johnson, 2000)) indicative of increased photoinhibition, i.e. PSII damage, under long-term fluctuating light treatments (Figure 2B).

**Figure 2.**
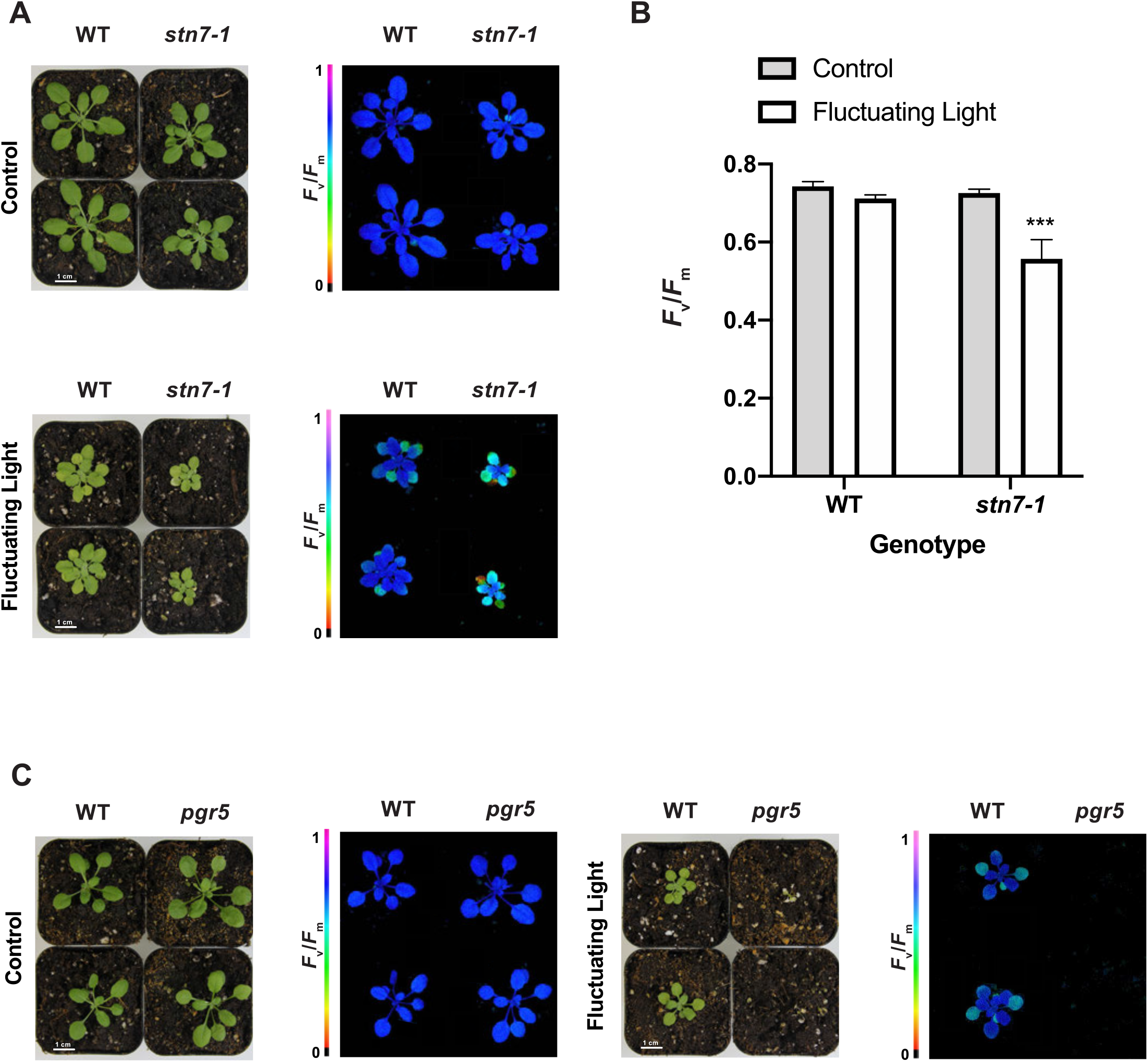
Arabidopsis WT and *stn7* phenotypes under constant light (control), and fluctuating light. A) Four week old plants which were exposed to constant light (90 μmol photons m^−2^ s^−1^) or fluctuating light (1 min at 900 μmol photons m^−2^ s^−1^ and 4 min at 90 μmol photons m^−2^ s^−1^) for the last two weeks. *stn7* plants under fluctuating light revealed decreased growth and *F*_v_/*F*_m_ values compared to WT under fluctuating light. B) Bar graph of mean *F*_v_/*F*_m_ (± SE, n=5). Asterisks indicate a statistically significant difference compared with WT (***P<0.0001, two-way ANOVA). C) Four-week-old plants exposed to fluctuating light. *pgr5* did not survive the treatment for more than 5 days compared to WT.

The extreme sensitivity of *pgr5* loss-of-function mutants to fluctuating light has been reported many times by independent groups (Suorsa et al., 2012; Thormählen et al., 2017; Trinh et al., 2019). The susceptibility is primarily attributed to a malfunctioning cyclic electron flow (CEF) cycle around PSI (Munekage et al., 2002). Therefore, *pgr5* was also tested in our newly developed low-cost growth setup. Because of the sensitivity to fluctuating light, *pgr5* and a set of WT plants were initially grown under constant light (12/12 h day-night cycles) for two weeks and then shifted from the lower shelf into the fluctuating light on the upper shelf. No *pgr5* mutant individual survived fluctuating light treatment longer than five days while all control plants under constant light conditions performed well (Figure 2C).

In summary, the data obtained show that our cost-effective fluctuating light plant growth rack delivers comparable results to previously published studies that used higher cost commercial solutions. The rack is easy to setup and, with costs below $650, represents a useful alternative for research groups with limited financial resources.

### Design of a sample holder kit for the IMAGING-PAM to improve throughput and data quality

The IMAGING-PAM can produce excellent images of chlorophyll fluorescence, but we found a few small additions to greatly improve the user experience by streamlining downstream analysis. The cost-effective plant growth racks described above enable more biological repetitions that include wild-type controls grown under both constant light and fluctuating light. To keep up with processing increasingly larger datasets, we reconfigured our IMAGING-PAM device to produce images with consistent plant placement and lighting conditions to facilitate more automation in the downstream analysis.

The sample holder kit includes a sample crate and standardized pot holder. First, a sample crate was built to accommodate nine of our 2” x 2” x 2⅛” (5 cm x 5 cm x 5.5 cm) pots (Figure 3A). The inner height of the crate was determined to ensure perfect camera focus at the lowest magnification. Second, holders for these nine pots (Figure 3B, Additional file 3) were milled using PVC (an alternative option is also for four 3” x 3” x 3.5” or 6.4 cm x 6.4 cm x 7.6 cm pots (Additional file 3)). A small notch was added to the upper right corner of the holders to allow for easy handling and consistent positioning of the plant holders even in the darkness when assaying dark-adapted plants. The height of the holders can be adjusted using the screws on each corner and should be fixed with a nut to fit the pots in the same vertical and horizontal position. All parts were made from standard PVC hard plastic, but other materials may be cheaper and perform equally well. However, it important to use opaque, low reflectance materials. All detailed technical schematics can be found in Additional file 3. Scientists working at institutions without machine shop are welcome to contact the corresponding author for assistance ordering through the Instrument Shop at WSU.

**Figure 3.**
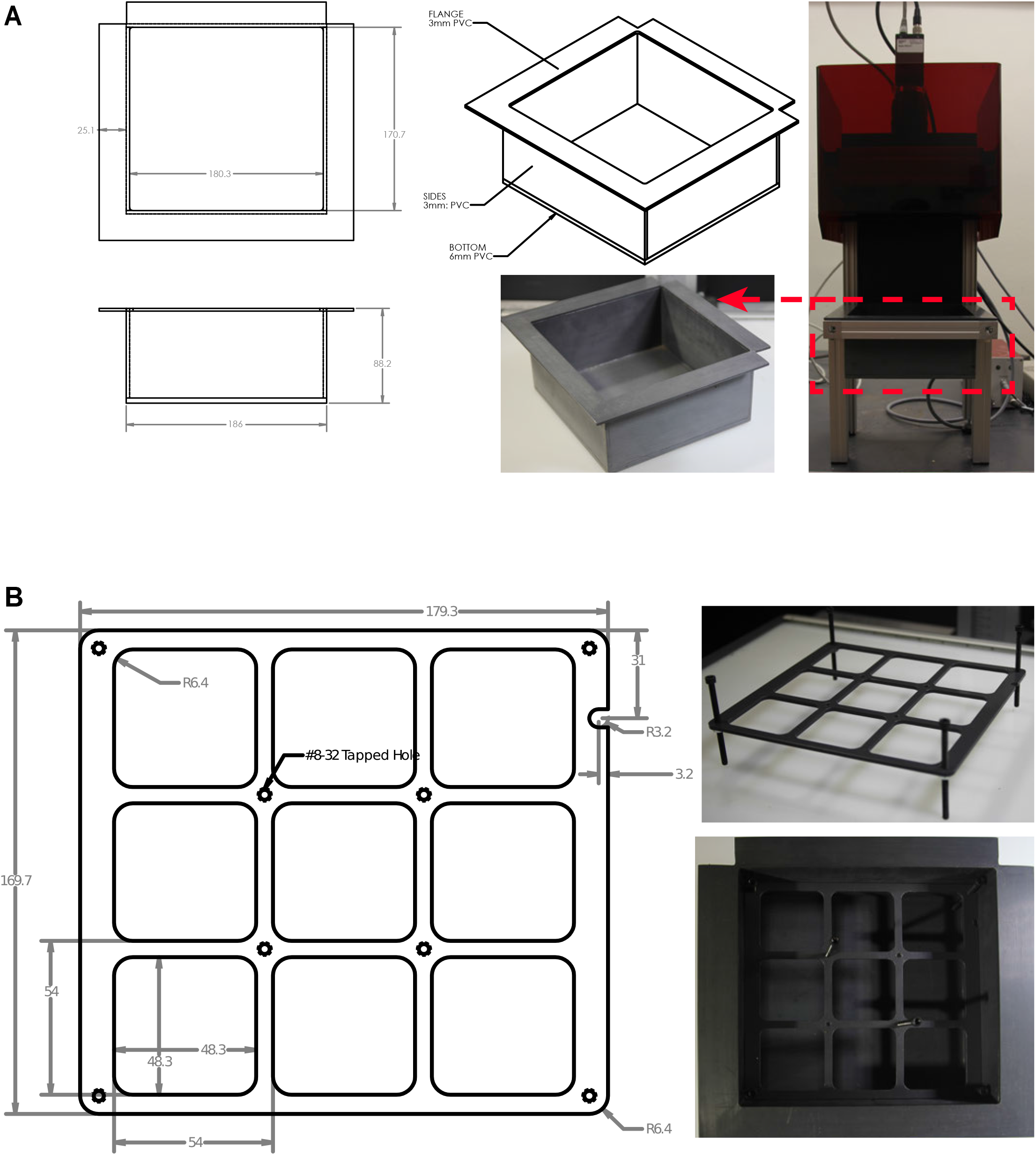
Reconfiguration of the Walz IMAGING-PAM. A) Drawing and image of the newly designed sample crate. B) Sample crate inserted in the IMAGING-PAM. C) Drawing and image of newly designed 9-pot-holders. Pot dimensions: 2” x 2” x 2⅛” (5 cm x 5 cm x 5.5 cm). The holders fit perfectly into the sample crate. The height of the holders can be adjusted with screws to ensure ideal pot-holder fit.

Although the working distance between the plants in the nine-plant pot holder and the camera lens is 2.6 cm longer than the 18.5 cm recommended by the manufacturer, this has no detectable effect of the image quality and the light pulse intensity. As shown in Figure 4, the reconfigured IMAGING-PAM delivers perfect plant images (*F*_v_/*F*_m_, NPQ shown in false colors) using *A. thaliana* wild-type plants vs. previously published *npq4-1* (Li et al., 2000) and *npq2-1* mutants (Niyogi et al., 1998) (21 days old, 12/12 h, constant light), with constitutive low NPQ and constitutively increased NPQ, respectively. Furthermore, the consistency of the setup, i.e. static position of the plants, is conducive for smooth time-lapse movies. This aids in visually tracking growth rates or phenotypic changes dependent on the plant developmental stage in specific mutant individuals. The holders ensure that each individual pot, and with that each individual plant, are recorded in the same position every time. The result is a much smoother time-lapse movie without the effect of plants bouncing around because of the difficulty of repositioning the plants in the same place for every measurement.

**Figure 4.**
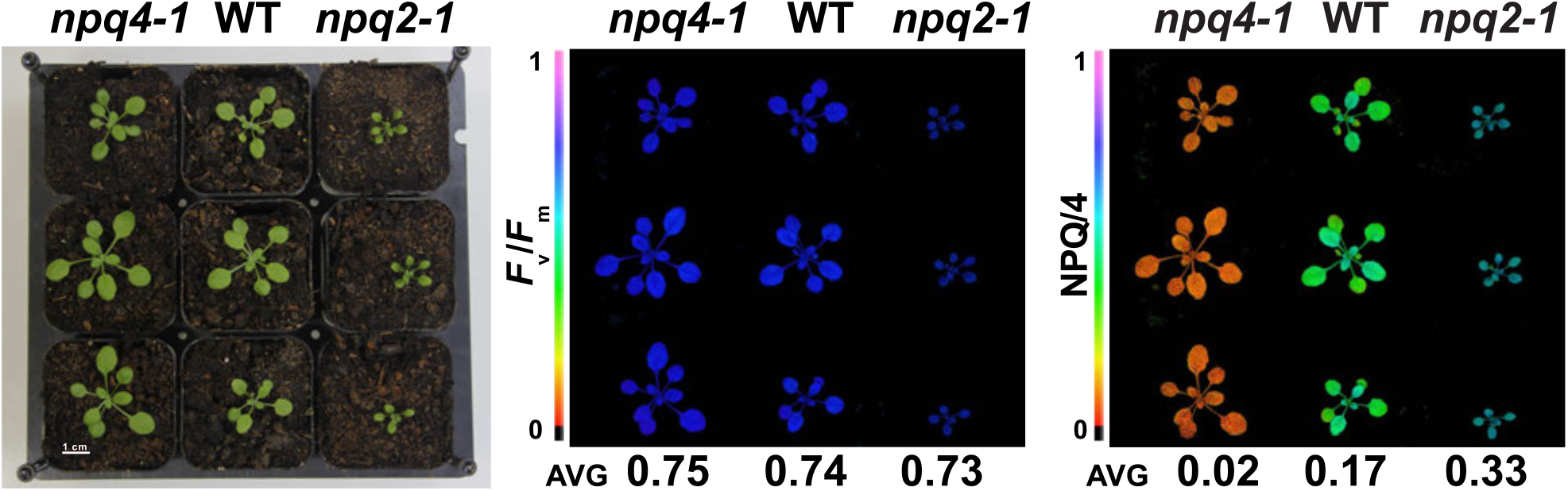
The reconfigured IMAGING-PAM with the newly designed sample crate and holders delivers perfectly focused false color images and values (*F*_v_/*F*_m_ and non-photochemical quenching NPQ/4) of four week old *npq4-1*, wild-type, and *npq2-1* plants grown in constant light (90 μmol photons m^−2^ s^−1^).

### Efficient analysis of images recorded with an IMAGING-PAM

The ImagingWinGigE freeware by Walz is useful to control the IMAGING-PAM camera. Additionally, its script function provides an option to run customized measurement protocols. However, the downstream analysis is cumbersome and time-consuming because each pim file (its native format) must be loaded separately and areas-of-interest (AOI, or region-of-interest ROI as it is commonly called) need to be manually assigned. The development of the sample crate and plant pot holder to fix the plant positions (Figure 3A-B) was largely motivated by the desire to automate the analysis of multiple files. Automation requires that sample plants always appear in the same location of an image, which our efforts described above accomplish as long as the camera settings are not changed.

We developed the ImagingPAMProcessing toolkit that includes scripts in Python and R to automate the phenotype extraction from a stack of measurement files and visualize the results. These scripts can be downloaded as a .zip via GitHub (https://github.com/CougPhenomics/ImagingPAMProcessing). The scripts in their current version feature: 1) automated plant recognition (leaf segmentation) in Python using PlantCV (Gehan et al., 2017). 2) automated genotype assignment from a separately provided metadata file 3) calculation of *F*_v_/*F*_m_, NPQ, YII (the Quantum yield of PSII), and plant surface area 4) false-color pictures to visualize heterogeneity 5) Rmarkdown report to visualize data quality and trends in the phenotypes 6) R script to create time-lapse videos of false-color pictures of each of the photosynthetic parameters.

### ImagingPAMProcessing Toolkit Setup

There are three main files that comprise the toolkit. The main script that processes the images is ProcessImages.py while postprocessingQC.Rmd and makeVideos.R facilitate visualizations. There are a few prerequisite steps before using the ImagingPAMProcessing toolkit:

1. The PIM files must be exported to a generic format, i.e. TIFF, which can be accomplished with the ImagingWinGigE software either manually (Figure 5) or by adding the “Export to Tiff File=” command at the end of running a custom ImagingWinGigE script. See diy_data/LemnaTec2.prg for an example. This results in a multi-frame TIFF file with the same structure as the PIM file. The filenames of the multi-frame TIFF files must be standardized with hyphens to uniquely identify each measurement protocol. For instance, in the example dataset: treatment (control or fluc), the date of measurement (formatted YYYYMMDD), and the sample id (tray #) to identify the files: fluc-20190901-tray2.tif
2. We use two configuration files, or metadata maps, to provide more information for downstream analysis. First, pimframes_map.csv contains the definition of each frame of the TIFF file and the corresponding induction period. The order of the frames is standardized from Walz and the first four frames will not change between protocols. The frames of the TIFF files are arranged such that frames one and two are Fo and Fm, respectively, and frames three and four Red Absorptivity and NIR Absorptivity, respectively. Additional frames come in pairs (five/six, seven/eight, etc) where each pair correspond to F’/Fm’ fluorescence measurements in the order they were captured. Note, if Fo and Fm were measured as the initial induction period, then these frames are repeated in frames five/six. There are 34 frames resulting from the default induction curve protocol accessed through the ImagingWin Induction Curve tab. Correspondingly, our pimframes_map.csv includes entries for frames 1-34, with 15 different induction periods (*F*_v_/*F*_m_ and 14 additional pairs of *F’/Fm*). The second configuration file is called genotype_map.csv and relates the genotype to the tray number and ROI position. An example of both files is provided in the GitHub repository.

**Figure 5.**
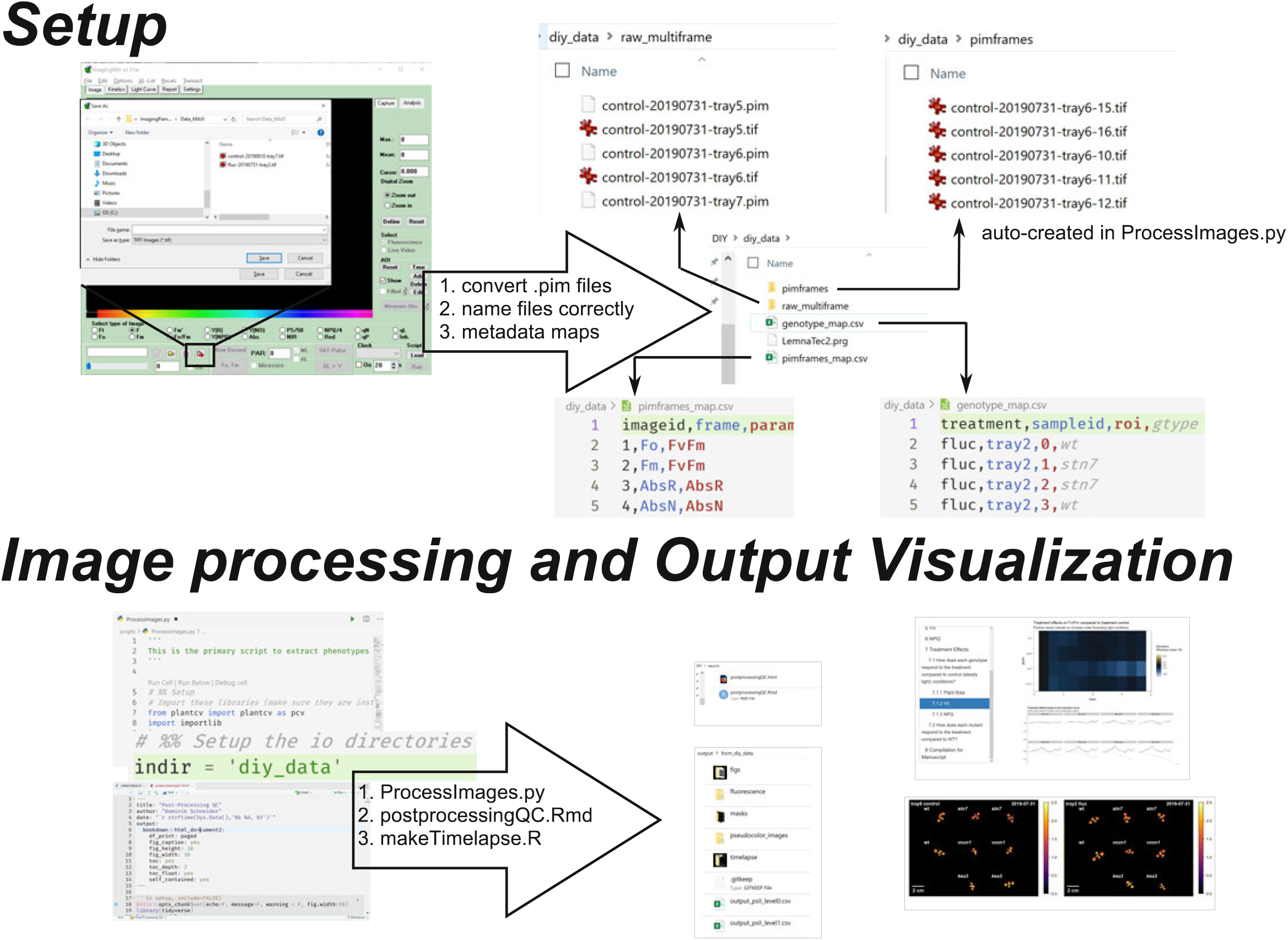
Schematic showing the critical steps to setup and run the scripts in the ImagingPAMProcessing toolkit.

### ProcessImages.py Customizations

In addition to the two configuration files, the user must update the `indir` variable in ProcessImages.py to point to their data directory. Additionally, there are three pieces of the image processing that may need to be adapted to the specific users’ imaging setup:

1. Image segmentation is generally quite specific to the imaging conditions. An automated estimate for the initial threshold value is provided based on Yen’s Algorithm (Yen et al., 1995), which is an entropy-based method implemented in the Python package scikit-image (van der Walt et al., 2014). This is followed by cleaning steps to remove small noise in the mask. In particular, we expect the cleaning steps found in src/segmentation/createmasks.py may need to be modified to adapt to unique imaging conditions from individual IMAGING-PAM setups. It should be noted that severe algae and moss growth due to overwatering will contaminate the images and make the image segmentation difficult. For more guidance on image segmentation we refer the reader to the excellent tutorials hosted by PlantCV (https://plantcv.readthedocs.io).
2. It is also likely the user will need to modify the locations of the ROIs to indicate where the plants are in the image. Even if using the 9 plant arrangement with the sample crate and 9 plant pot holders described in the text, it is likely the camera working distance will be slightly different and therefore the plant positions will be different relative to the image frame. In this case the location of the ROIs must be changed in the call to `pcv.roi.multi()` in scripts/ProcessImages.py. ROI coordinates can be adapted and visualized by stepping through the analysis with a single image with `pcv.params.debug = “plot”`. See the PlantCV documentation for details.
3. Our script outputs plant area that is automatically determined from the object detection algorithm implemented through PlantCV. It is important that each user updates the `pixel_resolution` variable for their own IMAGING-PAM setup to accurately convert pixels to mm^2^. This variable will be specific to the camera and working distance and can be found near the top of the main python script. This needs only be performed once as long as the camera settings remain constant. We recommend imaging a plant with a hole punch of a known size and then measuring the width in pixels of the hole using ImageJ. `pixel_resolution` is then calculated as diameter in mm of hole punch divided by diameter in pixels of hole punch.

### Post-Processing Report

In addition to the main python script for processing the image files, we also developed a report using RMarkdown (the source is found in the GitHub repository under reports/postprocessingQC.rmd) that can be compiled to html (Additional file 4) and is intended to provide a story-board-like overview of the extracted phenotypes. The user adjusts the variable `datadir` to point to the directory which contains the input images. Our first analysis shows whether all the data is present and if any of the QC flags were activated during image processing. In particular, we are interested in whether each plant was completely imaged and whether the plants remained independent in the image, i.e. did not overlap with each other at a given time point. A False value for each of these tests invalidates the results of the image processing and motivate the removal of these data points from further analysis. The next focus of the post-processing report is to visualize the trends in each phenotype for each plant. We plot timeseries of plant area, YII, and NPQ with bar plots and line plots because each plot type has unique advantages. Plotting using a prescribed pipeline makes it trivial to generate an array of figures quickly and simultaneously. Bulk visualization becomes important with more data being collected because it gives the researcher a starting point to identify the most interesting features of the data. It is also easy to identify data points that are out of range compared to the rest of a mutant panel. We find the RMarkdown report advantageous compared to separate plots because each section can be annotated and reads like a picture book. For example, in section 7 of our report, we are interested in the treatment effects. We clearly labeled the question we are interested in, can refer to the data manipulation used, and can evaluate multiple figures to address the questions. At the end we can compile any set of figures as required for publications (e.g. Figure 6).

**Figure 6.**
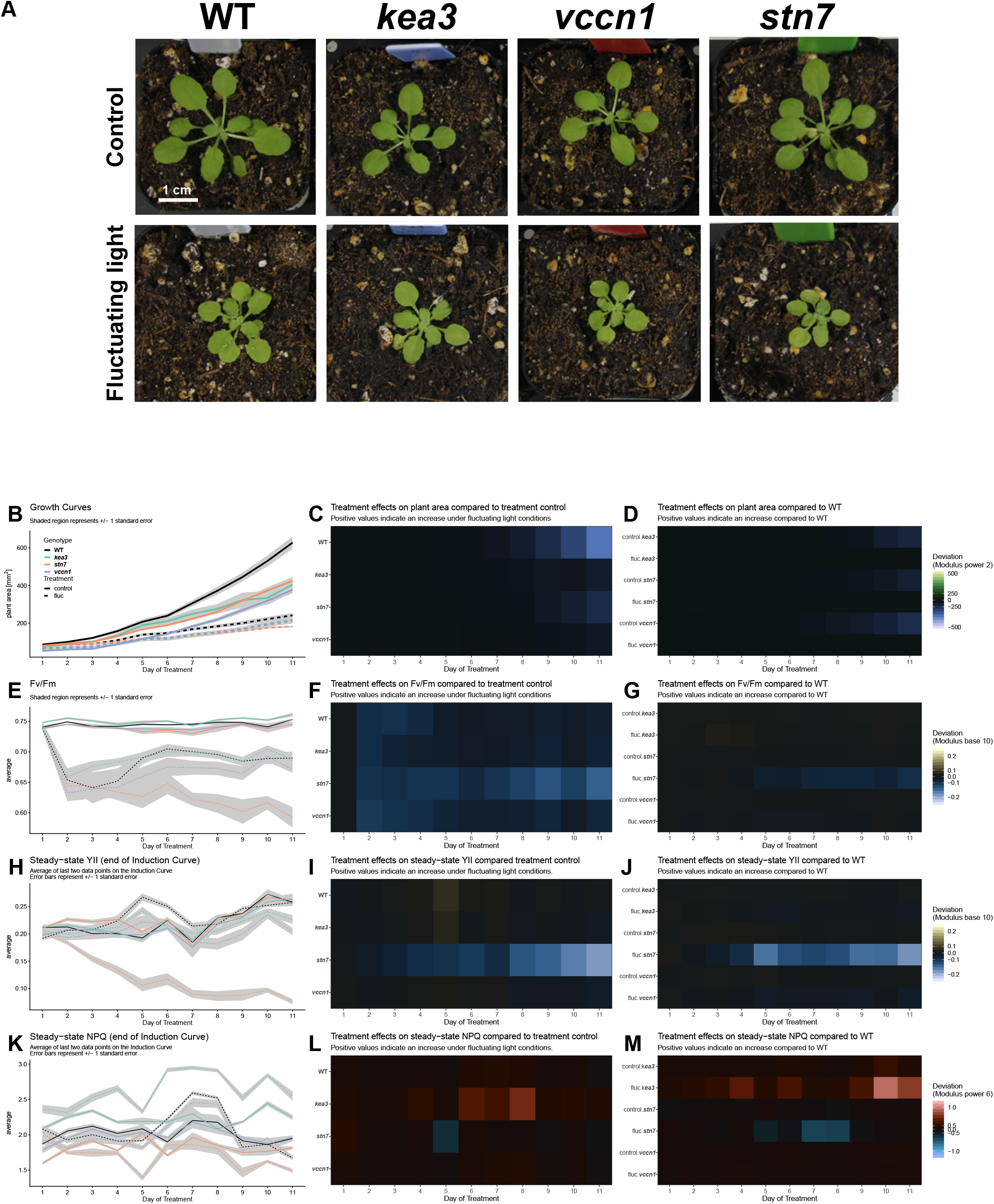
Data analysis from 11 day long phenotyping experiment. A) Four week old mutant lines and WT control plants after being subjected to constant light (control) or fluctuating light conditions. B-D) Growth behavior, E-G) *F*_v_/*F*_m_, H-J) YII, and K-M) NPQ throughout the experiment.

### Time-lapse Movies

Lastly, it is noteworthy that the ImagingPAMProcessing toolkit contains scripts/makeVideos.R which can compile *F*_v_/*F*_m_, YII, NPQ false color time-lapse movies into small-sized gifs which can be readily incorporated into slide presentations. The script automatically annotates plants with their genotype and creates a movie for each pair of trays. This script runs independently from the report. As mentioned earlier, the newly designed sample holder kit ensures that each individual plant is recorded in the same position every time. The resulting time-lapse movies of the sample dataset provided here can be found as Additional file 5.

### Testing the ImagingPAMProcessing toolkit using a diverse mutant panel recorded with the IMAGING-PAM

We employed the newly built growth rack (Figure 1) to record an eleven-day timeline of Arabidopsis loss-of-function mutants grown under two different light treatments to showcase the power and versatility of the ImagingPAMProcessing scripts. Specifically, we chose the *kea3* mutant which is affected in K^+^/H^+^ exchange across the chloroplast thylakoid membrane (Armbruster et al., 2014; Kunz et al., 2014) and the *vccn1/best1* mutant affected in thylakoid Cl^−^ ion flux (Duan et al., 2016; Herdean et al., 2016). Lastly, we added the previously mentioned *stn7* mutant which is compromised in its capability to adapt to changing light conditions (Figure 2A-B) (Bellafiore et al., 2005; Bonardi et al., 2005). The ion transport mutants served as points of reference as they were recently characterized in a five day dynamic environmental photosynthesis imager (DEPI) experiment (Cruz et al., 2016; Höhner et al., 2019). One half of the mutant panel was kept on the lower shelf of the plant growth rack, i.e. exposed to constant light (90 μmol photons m^−2^ s^−1^, 12/12 h day-night cycle) throughout its three-and-a-half-week life cycle. At an age of 14 days, the other half of plants was exposed to fluctuating light on the upper shelf (1 min at 900 μmol photons m^−2^ s^−1^, 4 min at 90 μmol photons m^−2^ s^−1^; 12/12 h day-night cycles). Data were recorded daily with the IMAGING-PAM for 11 days and plants photographed in true-color at the end of this period (Figure 6A). A single day of phenotyping alone yielded 1,448 data points (6 trays x 8 plants x 15 induction periods x 2 photosynthetic phenotypes + 48 estimates of plant area). The 11-day screening period resulted in 16,368 data points, and more phenotypes might be of interest in future experiments. Image standardization and a repeatable processing pipeline were critical to analyze and inspect results in a time-effective manner.

We used the ImagingPAMProcessing toolkit to estimate and visualize plant size and fitness. In doing so, it became obvious that the fluctuating light treatment adds a detrimental abiotic stress to all genotypes (Figure 6B-D). WT and all mutants lost about half of their biomass according to the surface area calculation our script performs. In general, WT plants always seemed to grow best. However, because our proof-of-concept dataset had only four plant individuals per genotype and per light treatment, we remain cautious to interpret any potential growth performance differences among genotypes within either treatment group.

Photosynthetic fitness was evaluated with *F*_v_/*F*_m_ and steady-state YII and NPQ. The *F*_v_/*F*_m_ plots revealed that only fluctuating light triggered genotype-specific *F*_v_/*F*_m_ changes over time. Initially, the onset of high light pulses damaged all genotypes (indicated by decreased *F*_v_/*F*_m_) for the first 4 days (Figure 6E). WT and *kea3* eventually recovered PSII function and from thereon revealed values slightly below those from the constant light control group. However, loss of *KEA3* seemed to have a protective effect on PSII, i.e. while the initial loss of *F*_v_/*F*_m_ on the first day in fluctuating light was equally strong as in WT, the recovery was faster such that *kea3* mutants reached equally high *F*_v_/*F*_m_ values but two days earlier than WT controls (Figure 6E-G). *F*_v_/*F*_m_ in *vccn1* mutants remained slightly below WT level, and *stn7* was clearly the most compromised mutant in our panel with continuously progressing PSII damage in the presence of fluctuating light throughout the experiment (Figure 6E-G).

In line with the documented damage to PSII (low *F*_v_/*F*_m_), steady-state YII also vanished dramatically in *stn7* treated with fluctuating light (Figure 6H-J). Under the same light treatment, the two mutants *kea3* and *vccn1* revealed only slightly diminished YII compared to WT controls (Figure 6H, J).

We investigated steady-state NPQ among mutants in response to light treatments (Figure 6K-M). Under constant light, only *kea3* showed slightly elevated NPQ compared to WT (Figure 6K, M). This matches earlier results at similar light intensities (Armbruster et al., 2016). NPQ for *stn7* mutants showed slightly depressed NPQ compared to WT whereas steady-state NPQ in *vccn1* mostly behaved like the wild-type control (Figure 6K, M), confirming recent results (Duan et al., 2016; Herdean et al., 2016). However, this situation changed when plants were treated with fluctuating light. The effect on steady-state NPQ in *kea3* and *stn7* mutant lines became strongly aggravated by fluctuating light in contrast to WT and *vccn1* (Figure 6K, L). In line with previous reports (Armbruster et al., 2016; Höhner et al., 2019), NPQ was noticeably increased in *kea3* compared to WT under the same conditions (Figure 6K, M) and compared to *kea3* mutants grown under constant light (control) (Figure 6K, L). The opposite effect was seen in the *stn7* mutant, where, in the presence of high light pulses, NPQ decreased compared to WT under the same conditions (Figure 6K, M) and compared to *stn7* mutants grown under control conditions of constant light (Figure 6K, L).

## Discussion

Over the last decade, plant science and photosynthesis research has made a big push towards gaining insights into complex physiological, biochemical, and genetic processes under more realistic growth conditions than the traditional lab regimes in which growth environments are kept as constant as possible (Vialet-Chabrand et al., 2017; Andersson et al., 2019). In this respect, light regimes represent a good example because light intensities in nature change frequently (Slattery et al., 2018). So far, we have only scratched the surface of understanding the traits responsible for the rapid cellular acclimation to these irregular challenges. Therefore, it is important to empower more scientists globally with cost-effective tools so everyone can apply more natural but reproducible growth conditions. The work presented herein shows that employing fluctuating light conditions in plant science does not required high-priced commercially built LED setups housed in climate chambers. As long as a dark space at constant room temperature is available, a simple setup made from online ordered parts delivers congruent results. By providing detailed instructions and the script to control the LED panels (according to the most commonly published fluctuating light conditions published) everyone interested should be able to quickly assemble the parts to apply the same experimental light conditions (Figure 1).

Using previously published mutants *stn7* and *pgr5* (Figure 2), we successfully validated our experimental setup by achieving similar results compared to past work (Grieco et al., 2012; Thormählen et al., 2017). As new fluctuating light susceptible mutants are isolated, it is important to compare them with both WT and mutants with known phenotypes under constant and fluctuating light in order to put the treatment effects in perspective. Our results provide confidence that experiments with our new plant growth racks will yield interesting and accurate phenotypes. A potential improvement to our design is to provide stronger background illumination as the 90 μmol photons m^−2^ s^−1^ is at the low end of the ideal *A. thaliana* light intensity range. Further, it would be advantageous to provide constant illumination closer to the average equivalent photon flux in the fluctuating light conditions which is 252 μmol photons m^−2^ s^−1^. The plant-to-light distance could be decreased to increase the photon flux in the constant light shelf at the expense of increased temperatures at the leaf level. Future experiments should evaluate the impact of this change.

Expanding experimental conditions and involving suitable, published genetic controls as a point of reference is good practice and highly advisable in light experiments. However, this also significantly expands the size of the experimental dataset and increases data analysis requirements. Employing automated phenotyping platforms with capabilities to record photosynthetic performance would be ideal but the high equipment costs can prevent access to phenotyping tools at most academic institutions. To cope with these challenges, we turned the most widely distributed camera-based chlorophyll fluorometer, the Walz IMAGING-PAM, into a semi-automated phenotyper with a few simple adjustments. A sample holder kit consisting of a crate and potholders (Figure 3) ensures that plants can be measured in the same spot even if moving the specimens in and out of a growth chamber. The slightly increased sample distance to the camera lens did not result in unfocused images or a detectable loss in measuring light intensities (Figure 4). All schematics can be found online to replicate our system or parts can be ordered through us (Additional file 3). Lastly, we also encourage users to maintain consistent timing of measurements to minimize differences due to duration of light exposure or circadian effects.

The minor positioning updates allowed us to design the ImagingPAMProcessing toolkit, a new open source analysis pipeline specifically designed for increasing the throughput of the Walz IMAGING-PAM. However, scientists could adapt our tools to rapidly analyze and plot large and complex experimental datasets from any fluorometer. The image processing scripts automatically attempts plant segmentation to distinguish between leaf and background using the open source PlantCV phenotyping toolbox (Gehan et al., 2017). Common photosynthetic phenotypes and plant area are extracted per plant and can be visualized and analyzed in relation to treatment, time, and genotype. We specifically focus on highlighting differences between a genotype control and a treatment control and provide the ability to create time-lapse movies of each phenotype for each plant.

To validate the script and to provide interested users with a training dataset, we recorded an eleven-day fluctuating light experiment using mostly genotypes recently tested in a five day long Dynamic Environmental Photosynthetic Imaging run (Höhner et al., 2019) (Figure 6). In line with earlier studies, we found that all genotypes were affected by fluctuating light (Vialet-Chabrand et al., 2017; Schneider et al., 2019). Leaf surface area in WT plants decreased by more than half. As reported before, we also saw evidence that growth of *stn7* mutants was especially impacted by fluctuating light which triggered dramatic decreases in *F*_v_/*F*_m_ and YII (Tikkanen et al., 2010; Grieco et al., 2012). Our observations of steady-state NPQ and YII in thylakoid ion transport mutants *kea3* and *vccn1* are also in line with other recent reports of these mutants (Dukic et al., 2019; Höhner et al., 2019).

## Conclusions

Fluctuating growth light conditions represent a cornerstone in understanding acclimation processes in photoautotrophic organisms. We have shown that high priced LED climate chambers and phenotyping equipment are not necessarily required to unveil the underlying genes involved in light stress acclimation processes. The simple construction of our micro-controller-based LED light racks and minor hardware modifications to the IMAGING PAM allow the application of our newly developed ImagingPAMProcessing toolkit. The wealth of data collected and analyzed in this way can provide new and highly useful insights. The tools introduced here are not limited to plant science but will also help to streamline genetic screens and physiology experiments in algae and cyanobacteria. For instance, the use of micro-multiwell plates in fixed positions in the IMAGING-PAM should allow for straight-forward application of the ImagingPAMProcessing toolkit. Accordingly, we encourage others to pick up the open source toolkit and adapt and expand it with new features.

## Methods

### Plant growth conditions

Wild type (WT) *Arabidopsis thaliana* accession Columbia-0 (Col-0) and mutant seeds were EtOH surface sterilized, stratified for two days at 4°C, and grown on ½ Murashige & Skoog (MS) 1 % (w/v) phytoagar plates pH 5.8 for one week at 90 μmol photons m^−2^ s^−1^ constant illumination in a 12/12 h day-night cycle at 22°C. At an age of seven days, seedlings designated for constant light conditions were potted into 2” x 2” x 2⅛” pots (Item #: 1665 by Anderson Pots, Portland, OR, USA) and grown under the same light condition until the end of their life cycle.

If individuals were designated for fluctuating light treatment, plants were initially grown for two weeks in constant light (90 μmol photons m^−2^ s^−1^) and then moved into fluctuating light (1 min at 900 μmol photons m^−2^ s^−1^ and 4 min at 90 μmol photons m^−2^ s^−1^ for two weeks.

Light intensities were careful monitored using a MQ-200 Quantum Separate Sensor with Handheld Meter and a data logger (Apogee Instruments, Inc. Logan, UT, USA). Both the LED grow lights and 1500W LED produce broad spectrum light from blue to infra-red with wavelengths ranging from 400 nm to 760 nm, similar to the sun. Their technical specifications can be found at https://www.suncolighting.com/pages/manuals-downloads and https://www.amazon.com/HIGROW-Double-Spectrum-Greenhouse-Hydroponic/dp/B075QGZKD2, respectively.

### Plant mutant isolation and information

The *vccn1-1* (SALK_103612) T-DNA insertion line (Herdean et al., 2016) was ordered from the ABRC stock center. Homozygous individuals were isolated through PCR-based genotyping using the WT primer combination: *VCCN1* 5’ UTR fwd (5’-3’: catgtcatgtgaagtgaagtgaag)/*VCCN1* rev (GCTGCAATGTAACGAAGAAGC) yielding a 1129 bps product and the KO primer combination *VCCN1* 5’ UTR fwd (5’-3’: catgtcatgtgaagtgaagtgaag)/ Salk LBb1.3 (5’-3’: attttgccgatttcggaac) to produce a ~ 500 bps product.

### Accession Numbers for this study

Additionally, the following homozygous loss-of-function mutant lines were employed in this study: *npq4-1* (Li et al., 2000), *npq2-1* aka *aba1-6* (CS3772, Niyogi et al., 1998), *kea3-1* (Gabi_170G09; Armbruster et al., 2014), *stn7-1* (SALK_073254, (Bellafiore et al., 2005; Bonardi et al., 2005)), *pgr5-1* (Munekage et al., 2001).

### Pulse-Amplitude-Modulation (PAM) fluorescence spectroscopy

A MAXI version IMAGING-PAM (IMAG-K7 by Walz GmbH, Effeltrich, Germany) was employed in all experiments were photosynthesis-related parameters were recorded. Before each measurements, plants were positioned in the newly designed plant holders. Subsequently, plants were dark-adapted for 15 minutes followed by recording of a standard induction curve at 186 μmol photons m^−2^ s^−1^ actinic light. All data were analyzed with the new ProcessImages.py script and for comparison also using the ImagingWinGigE freeware by Walz.

## Declarations

### Ethics approval and consent to participate

Not applicable

### Consent for publication

Not applicable

### Availability of data and materials

The scripts described in the text can be downloaded from https://github.com/CougPhenomics/ImagingPAMProcessing and the accompanying 11 day dataset can be downloaded from https://doi.org/10.17605/OSF.IO/P32AY

### Authors’ contributions

HHK designed the research and the sample holder kit, built the fluctuating light growth racks, and wrote the manuscript. DS programed the PSII processing pipeline, coded the R data potting toolkit, analyzed experimental data, designed figures, wrote sections of the manuscript, and edited the manuscript. LSL carried out all plant experiments, analyzed experimental data, and designed figures. JDC wrote the micro-controller script. ML and HK isolated *vccn1-1* line and edited the manuscript. All authors read and approved the final manuscript.

## Acknowledgements

We thank Dave Savage from the Technical Services Instrument shop at WSU for help with the design of the sample holder kit. We are very grateful for advice on pim file conversion provided by the scientists at Walz. Furthermore, we thank Dr. Ute Armbruster at MPI Golm for providing *pgr5* mutant seeds. HHK thanks The Helen Riaboff Whiteley Center and the people at the UW Friday Harbor Labs for providing a writing refuge.

## Additional Files

Additional File 1.

A) Background and fluctuating light mode of the growth racks. B) Extension or single unit fluctuating light shelf. Shown are both operation modes.

Additional File 2.

Fluctuating light script to flash the Adafruit (Arduino-type) micro-controller.

Additional File 3.

Schematics of the sample holder kit. Sample crate and plant pot holders are compatible with the IMAGING PAM.

Additional File 4.

Post-processing quality control output generated by the ImagingPAMProcessing toolkit. The proof-of-concept dataset from this was used for this output example.

Additional File 5.

Time-lapse movies for *F*_v_/*F*_m_, steady-state YII, and steady-state NPQ generated from the proof-of-concept dataset.

