## Additional File1 for "Fluctuating light experiments and semi-automated plant phenotyping enabled by self-built growth racks and simple upgrades to the IMAGING-PAM"

### Additional File 1

**A** Extension or single unit Fluctuating Light Shelf

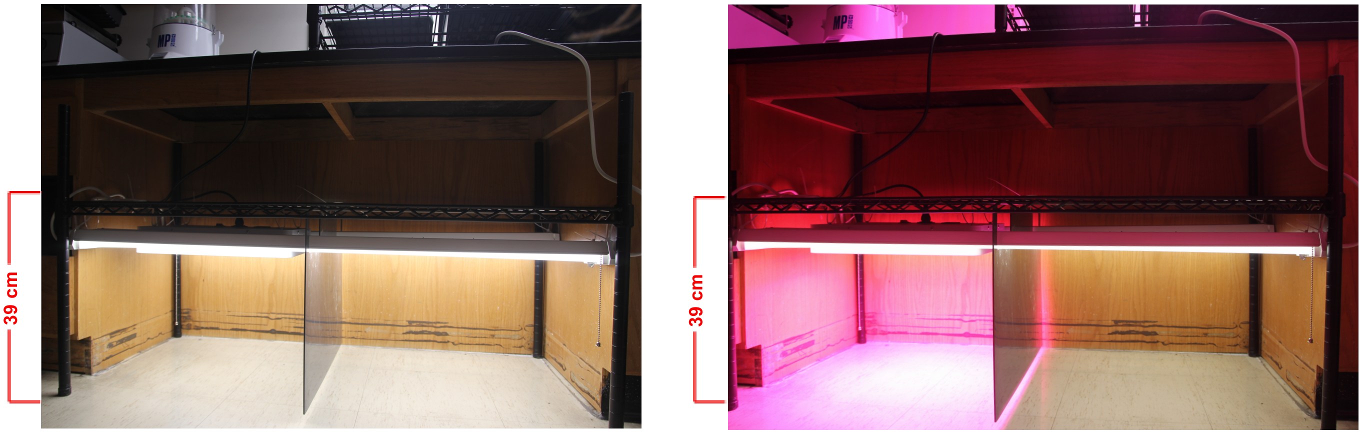


Fluctuating Light    Constant Light    Fluctuating Light    Constant Light

**B** Full Fluctuating Light Growth Rack

Background Light Mode    Fluctuating Light Mode

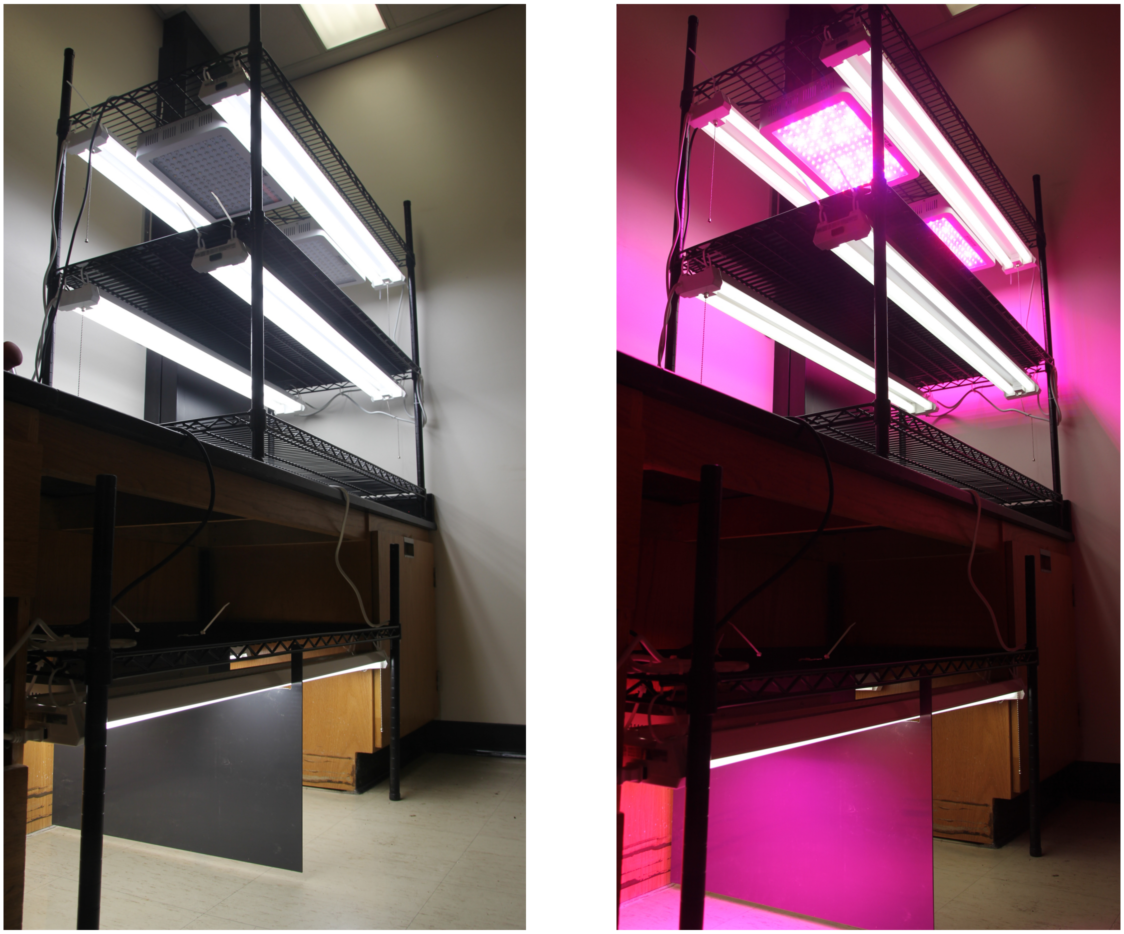
