## Additional File2 for "Fluctuating light experiments and semi-automated plant phenotyping enabled by self-built growth racks and simple upgrades to the IMAGING-PAM"

/*
 

Arduino script that controls the relay that turns power on and off to the grow light.

 */

// These are constants that won't change. Used here to see a pin number : 
const int ledPin = 13;     // the number of the pin that supplies power to the relay switch that controls the growlight power.

//Time constants that control the length of time on or off. Note they are in milliseconds. You can change this for light on off time. 
const long lighton = 60000; //milliseconds; equivalent to 1 min
const long lightoff = 240000; //milliseconds; equivelent to 4 min

// the setup function runs once when you press reset or power the microcontroller. The external light timer controls power to the board throughout the day
void setup() {
 // initialize digital pin 13 as an output. This controls the relay switch which in turn controls the grow light being on or off. 
 pinMode(ledPin, OUTPUT); 
}

// the loop function runs over and over again forever until power to the microcontroller is turned off by the outlet timer 
void loop() { 
 digitalWrite(ledPin, HIGH); // turn the growlight on (HIGH is the voltage level)
 delay(lighton);            // wait for lighton time.
digitalWrite(ledPin, LOW);   // turn the growlight off by making the voltage LOW
delay(lightoff);             // wait for lightoff time
}
