## Additional File3 for "Fluctuating light experiments and semi-automated plant phenotyping enabled by self-built growth racks and simple upgrades to the IMAGING-PAM"

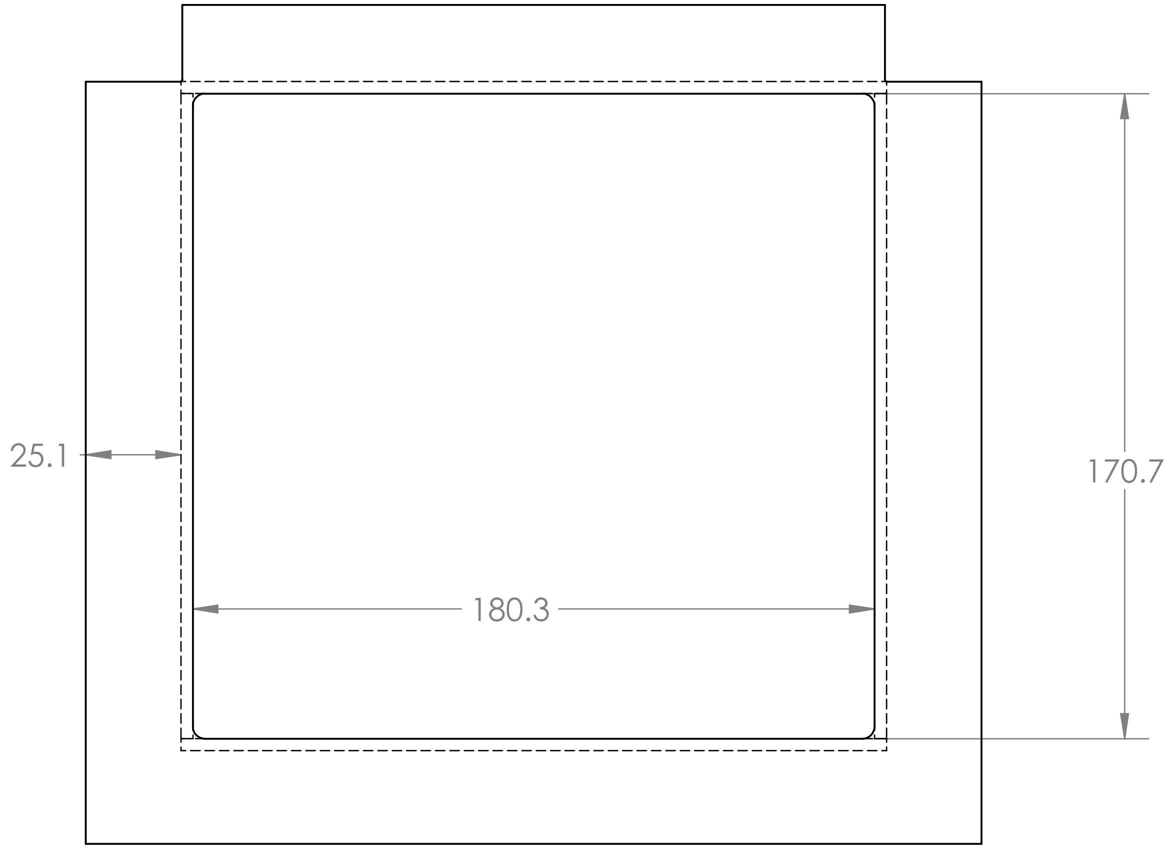


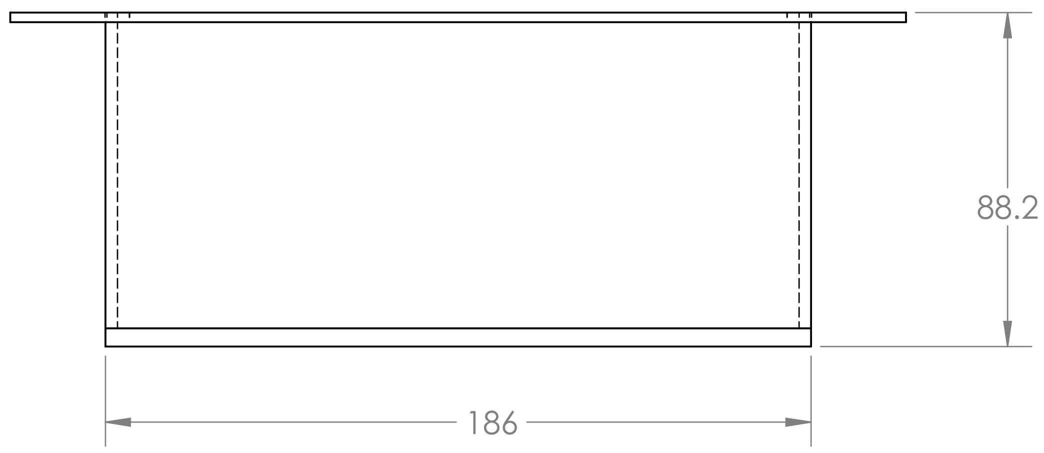


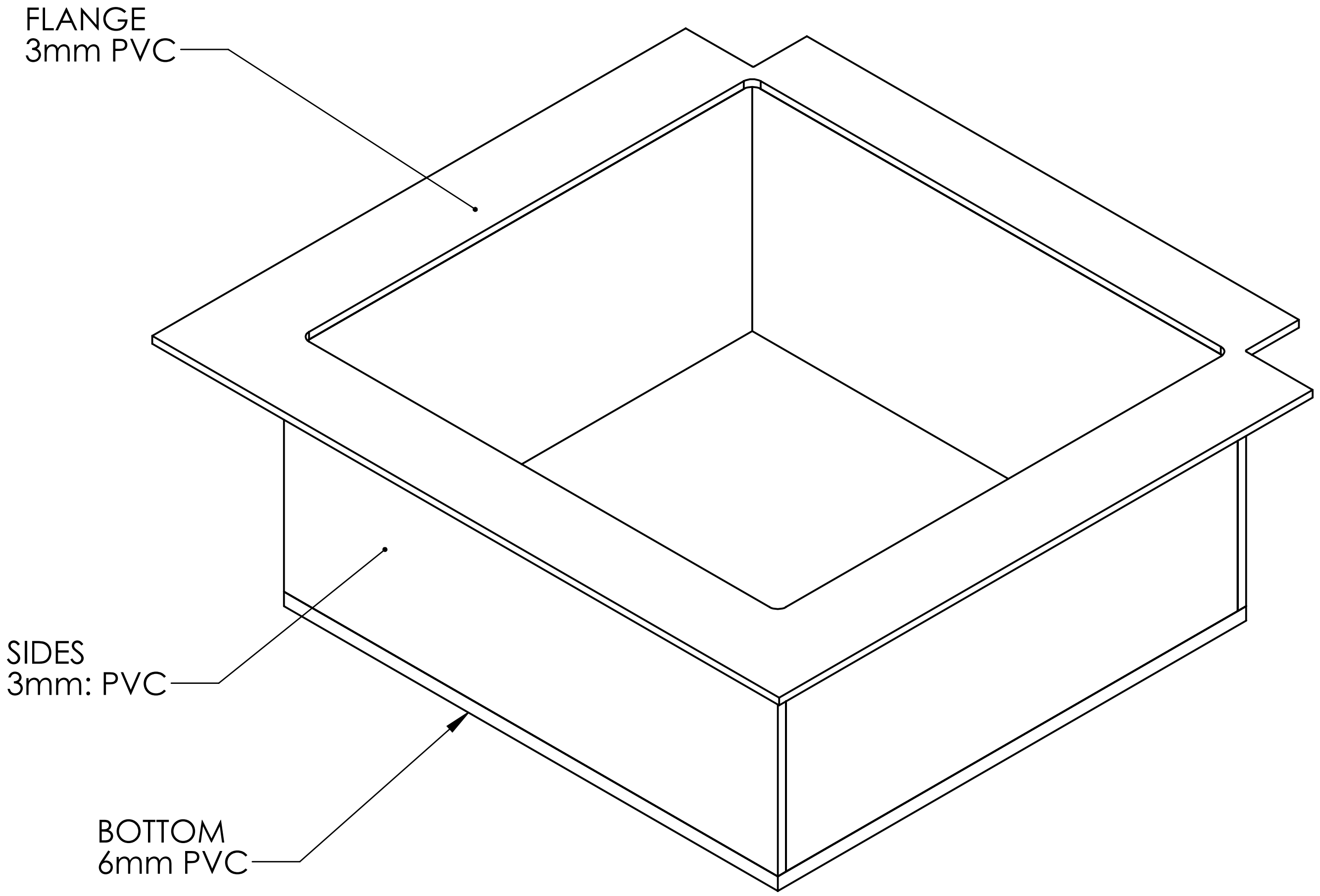


SOLVENT WELDED ASSEMBLY


THIS DRAWING IS PROPERTY OF WASHINGTON STATE UNIVERSITY DEPARTMENT OF TECHNICAL SERVICES.
ANY REPRODUCTION OR FABRICATION PROHIBITED WITHOUT WRITTEN PERMISSION OF THE OWNER.


UNLESS OTHERWISE SPECIFIED:

DIMENSIONS ARE IN MM
TOLERANCES:
FRACTIONAL ± 1/64
ANGULAR: MACH ± .1° BEND ±

MATERIAL AS NOTED

FINISH

DO NOT SCALE DRAWING

| | |
|---|---|
| DRAWN | SAVAGE |
| DATE | 6/5/19 |
| SCALE | 1:2 |
| SHEET | |
| JOB # | 430B/19 |

COMMENTS:

FILEPATH:

WSU TECHNICAL SERVICES

TITLE:

PLANT PHOTO CRATE

| SIZE | DWG. NO. | REV |
|---|---|---|
| B | | |

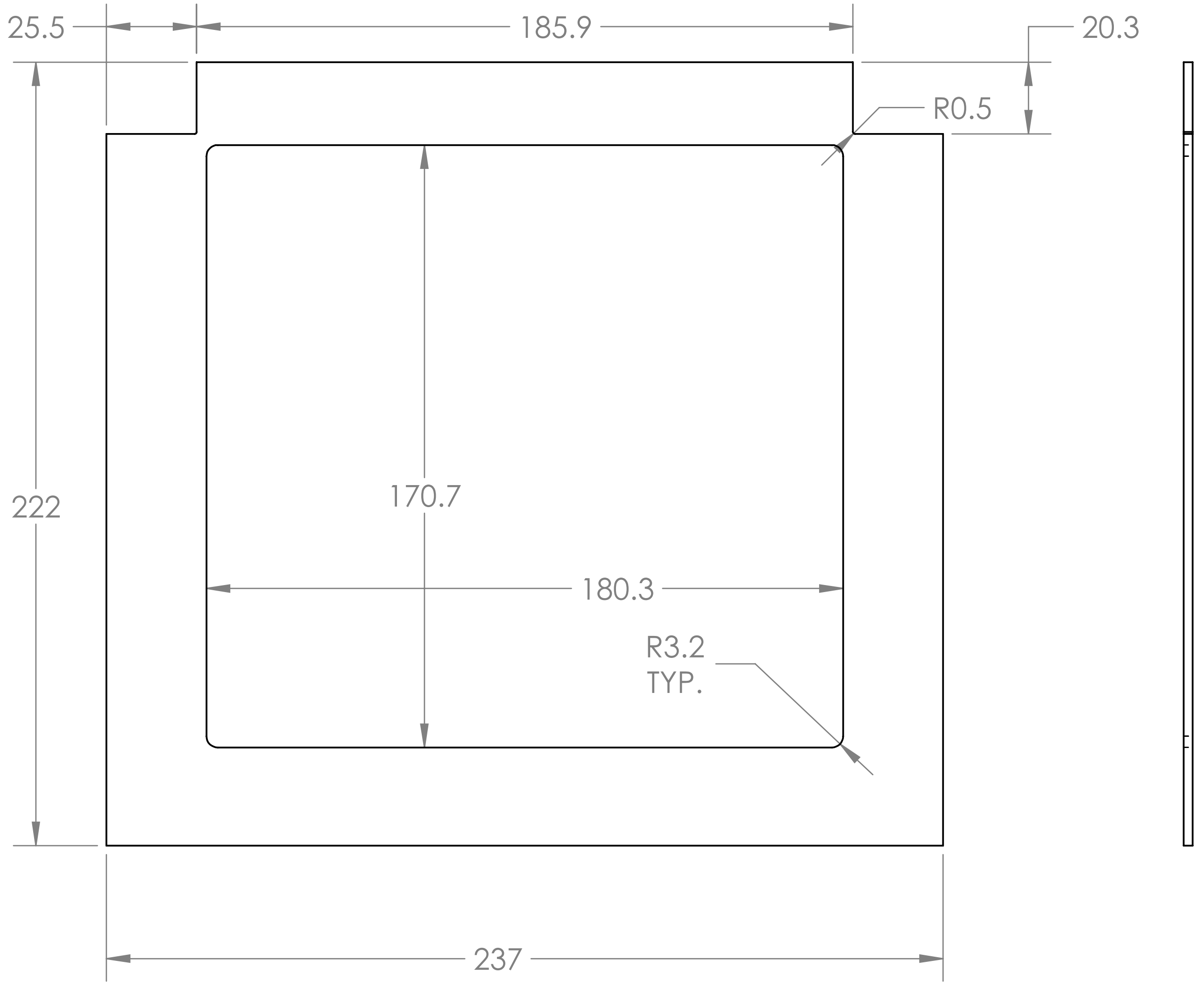


FLANGE:
3MM PVC

| UNLESS OTHERWISE SPECIFIED: | | | WSU TECHNICAL SERVICES |
|---|---|---|---|
| DIMENSIONS ARE IN MM<br>TOLERANCES:<br>FRACTIONAL ± 1/64<br>ANGULAR: MACH ± .1° BEND ± | DRAWN | SAVAGE | TITLE: |
| | DATE | 6/5/19 | PLANT PHOTO CRATE |
| | SCALE | 1:2 | |
| | SHEET | | |
| | JOB # | 430B/19 | |
| MATERIAL AS NOTED | COMMENTS: | | SIZE B · DWG. NO. · REV |
| FINISH | | | |
| DO NOT SCALE DRAWING | FILEPATH: | | |

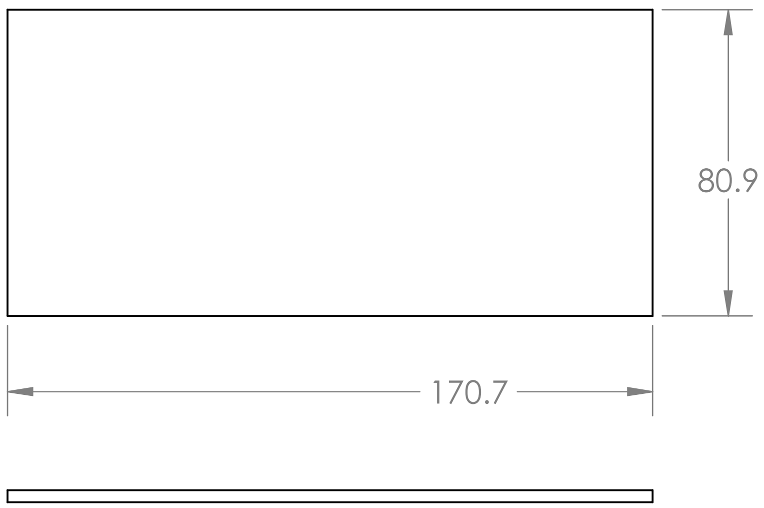


SIDES
QNT'Y: 2
MAT'L: 3MM PVC

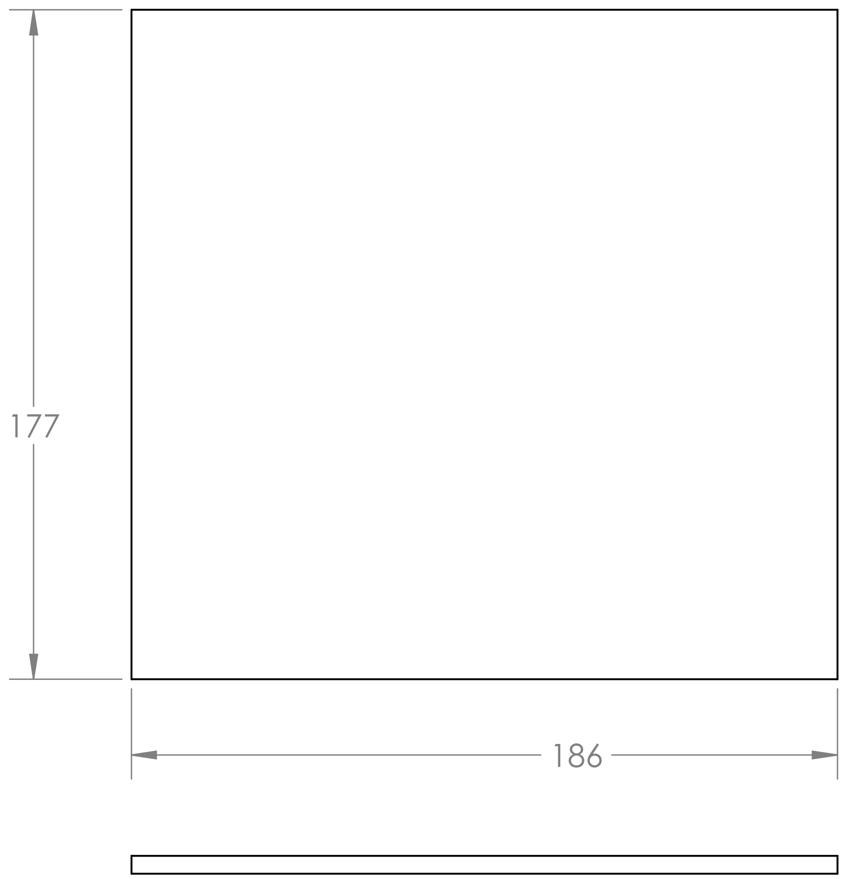


BOTTOM
MAT'L: 6MM PVC

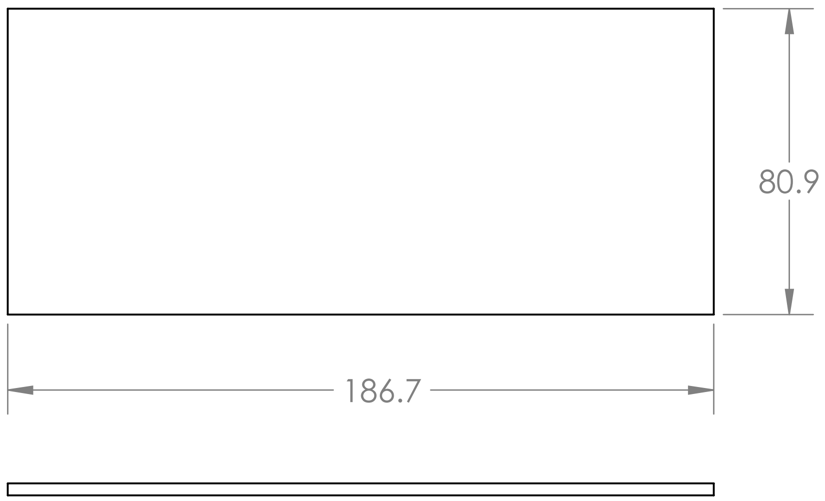


FRONT AND BACK
QNT'Y: 2
MAT'L: 3MM PVC

| | | | |
|---|---|---|---|
| UNLESS OTHERWISE SPECIFIED: DIMENSIONS ARE IN MM TOLERANCES: FRACTIONAL ± 1/64 ANGULAR: MACH ± .1° BEND ± | DRAWN | SAVAGE | WSU TECHNICAL SERVICES |
| | DATE | 6/5/19 | TITLE: PLANT PHOTO CRATE |
| | SCALE | 1:2 | |
| | SHEET | | |
| | JOB # | 430B/19 | |
| MATERIAL AS NOTED | COMMENTS: | | SIZE B / DWG. NO. / REV |
| FINISH | | | |
| DO NOT SCALE DRAWING | FILEPATH: | | |

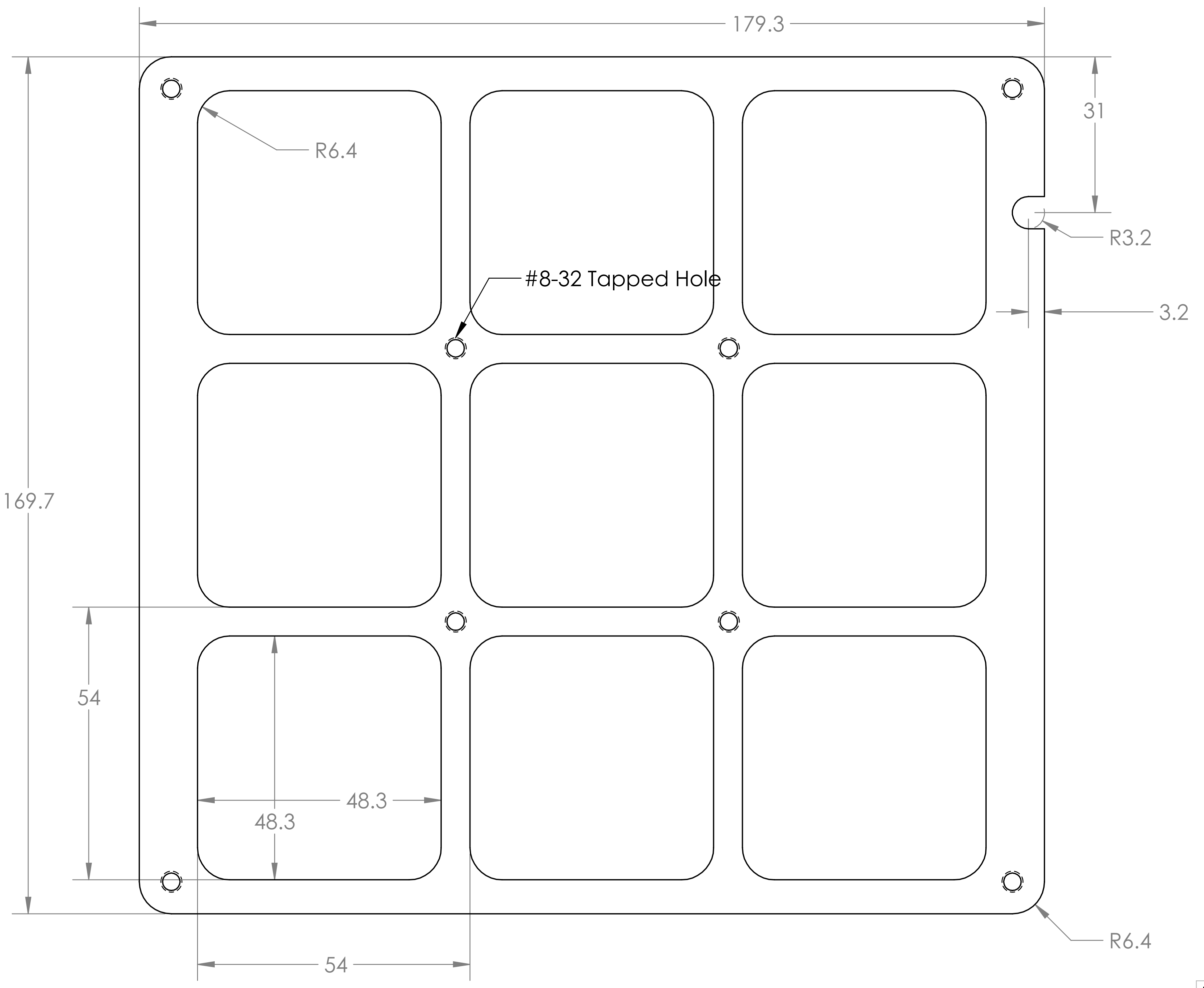


2-INCH POT

UNLESS OTHERWISE SPECIFIED:

DIMENSIONS ARE IN INCHES
TOLERANCES:
FRACTIONAL ± 1/64
ANGULAR: MACH ± .1° BEND ±
TWO PLACE DECIMAL ± .02
THREE PLACE DECIMAL ± .002

MATERIAL
3MM PVC

FINISH

DO NOT SCALE DRAWING

| | |
|---|---|
| DRAWN | SAVAGE |
| DATE | 7/23/19 |
| SCALE | 1:1 |
| SHEET | |
| JOB # | 430B/19 |

COMMENTS:

FILEPATH:

WSU TECHNICAL SERVICES

TITLE:

PLANT PHOTO CRATE

| SIZE | DWG. NO. | REV |
|---|---|---|
| B | | |
