## Additional File4 for "Fluctuating light experiments and semi-automated plant phenotyping enabled by self-built growth racks and simple upgrades to the IMAGING-PAM"

Post-Processing QC


### Post-Processing QC

###### Dominik Schneider

###### Sep 20, 2019

### 1 Read data from image analysis

### 2 Check image results

During image processing the detected objects are checked for:

1. whether the plant is completely in frame (top row)
2. whether the plants are unique (bottom row)

### 3 Subset Valid data

For subsequent plots we will remove invalid datapoints based on the quality checks above. These are saved as the “level1” dataset.

### 4 Plant Area

### 5 YII

#### 5.1 Steady-state YII

### 6 NPQ

#### 6.1 Steady-state NPQ

### 7 Treatment Effects

#### 7.1 How does each genotype respond to the treatment compared to control (steady light) conditions?

##### 7.1.1 Plant Area

##### 7.1.2 YII

###### 7.1.2.1 Steady-state YII

##### 7.1.3 NPQ

###### 7.1.3.1 Steady-state NPQ

#### 7.2 How does each mutant respond to the treatment compared to WT?

##### 7.2.1 Plant Area

##### 7.2.2 YII

###### 7.2.2.1 Steady-state YII

##### 7.2.3 NPQ

###### 7.2.3.1 Steady-state NPQ

### 8 Compilation for Manuscript
