## Supplementary figures and images for "Fluctuating light experiments and semi-automated plant phenotyping enabled by self-built growth racks and simple upgrades to the IMAGING-PAM"

### FvFm_YII_tray5_x_tray2.gif

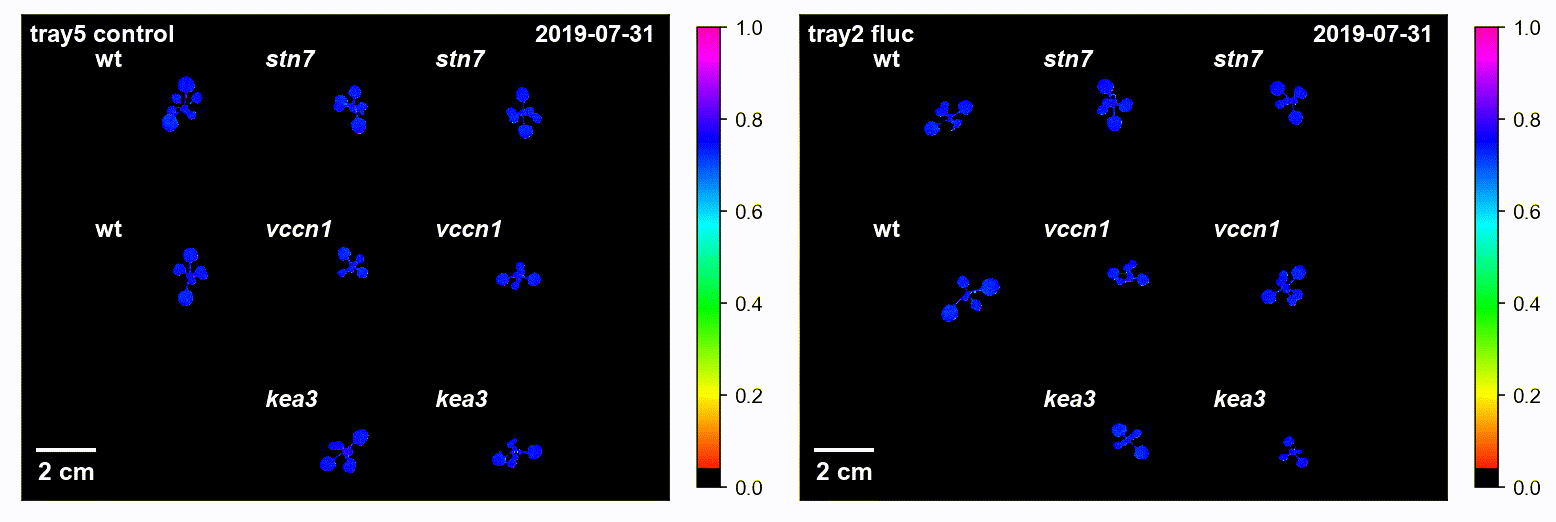

### FvFm_YII_tray5_x_tray3.gif

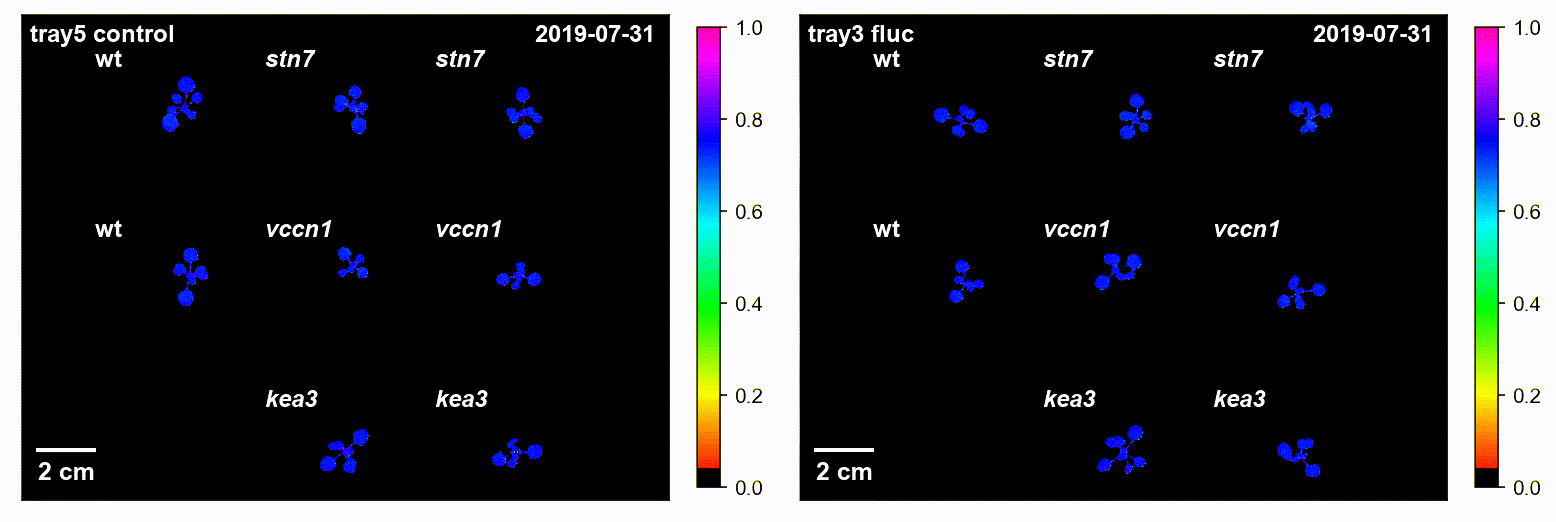

### FvFm_YII_tray5_x_tray4.gif

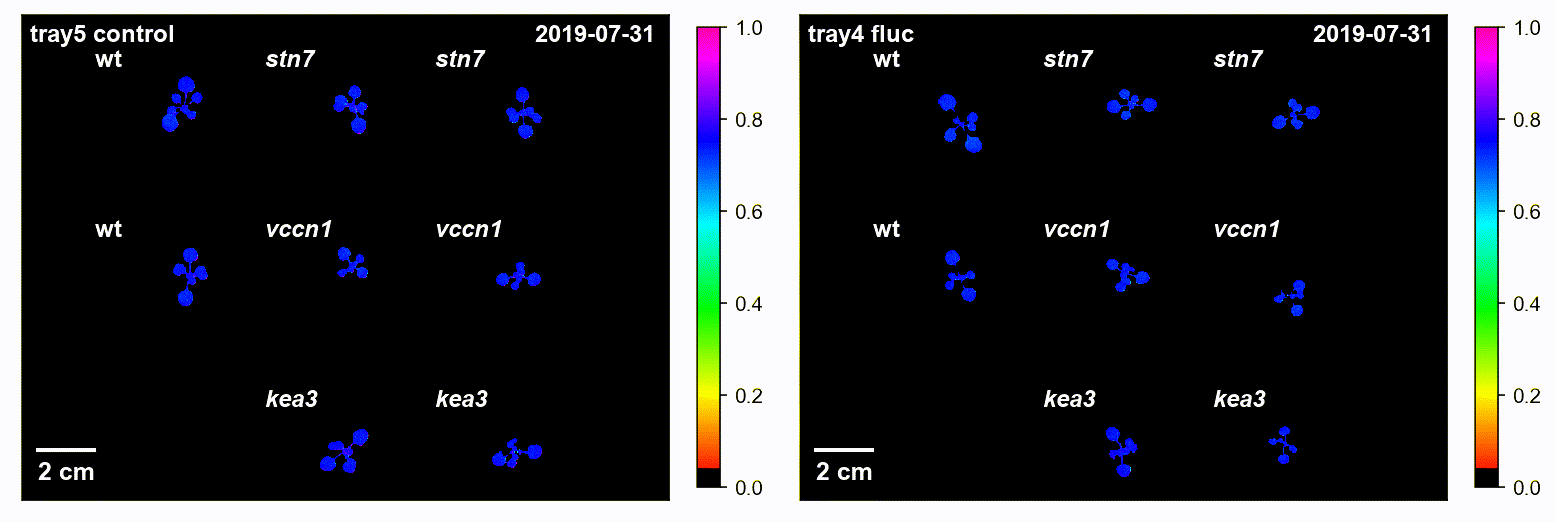

### FvFm_YII_tray6_x_tray2.gif

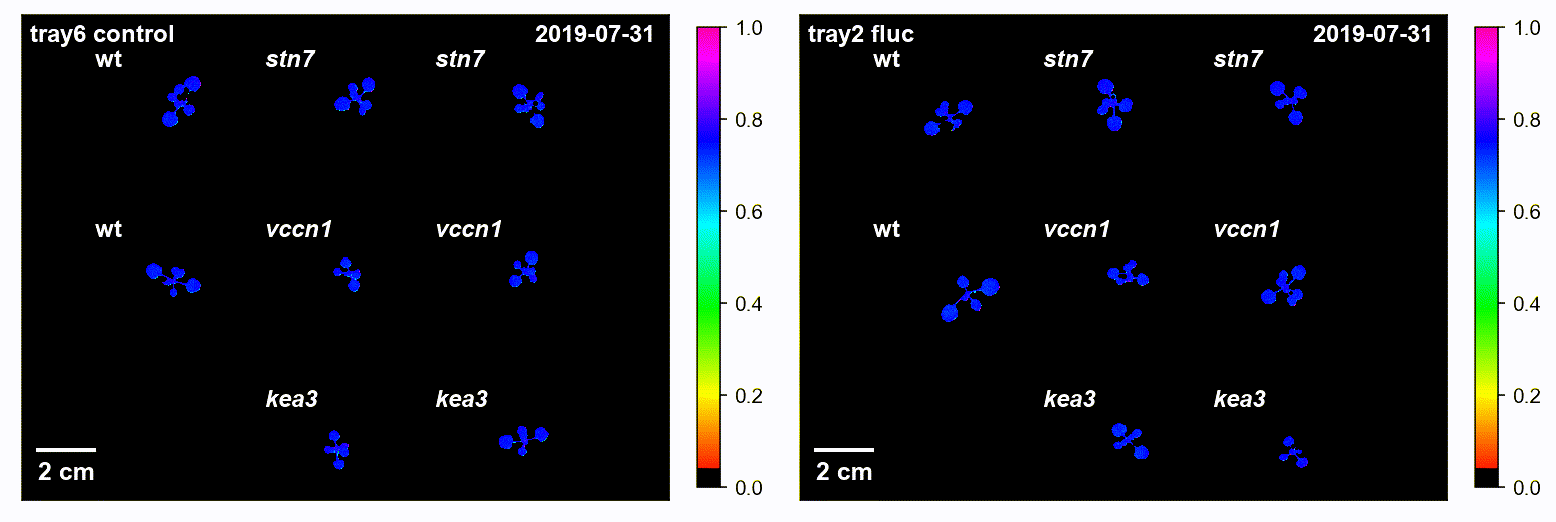

### FvFm_YII_tray6_x_tray3.gif

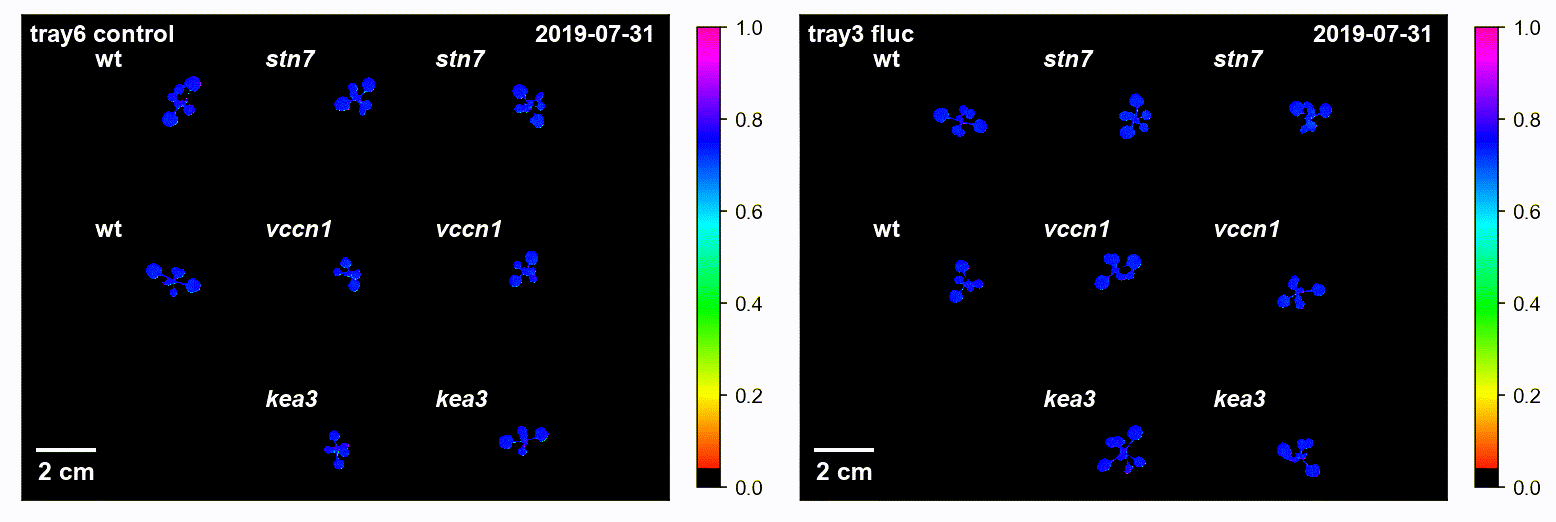

### FvFm_YII_tray6_x_tray4.gif

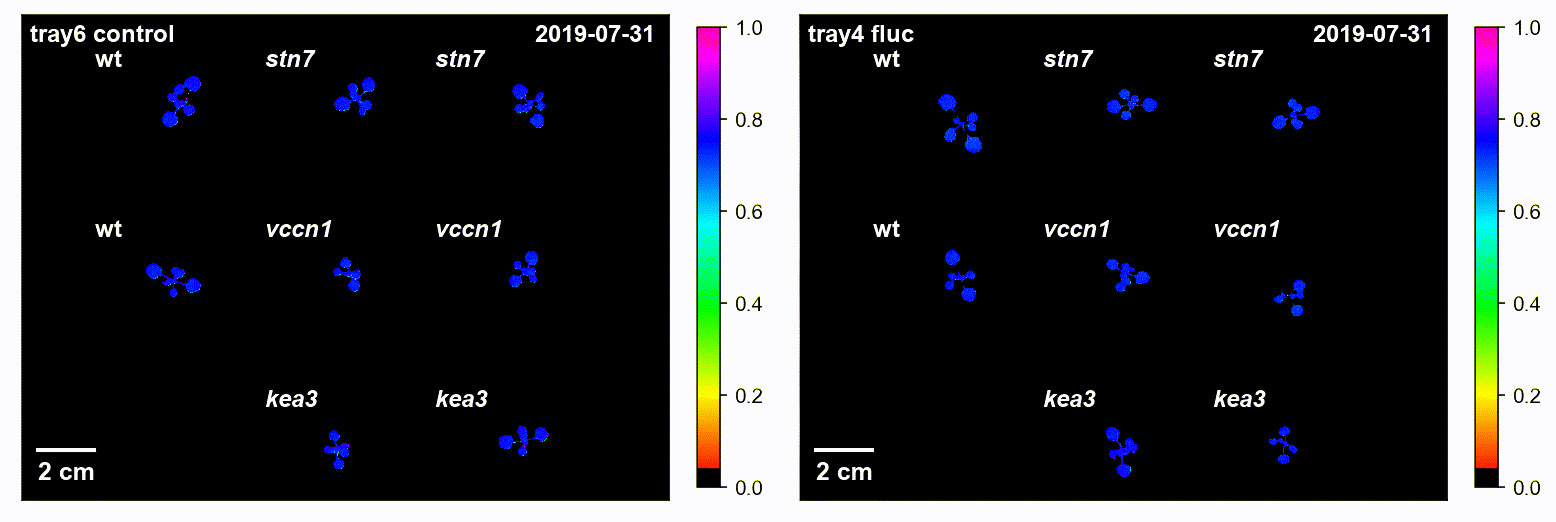

### FvFm_YII_tray7_x_tray2.gif

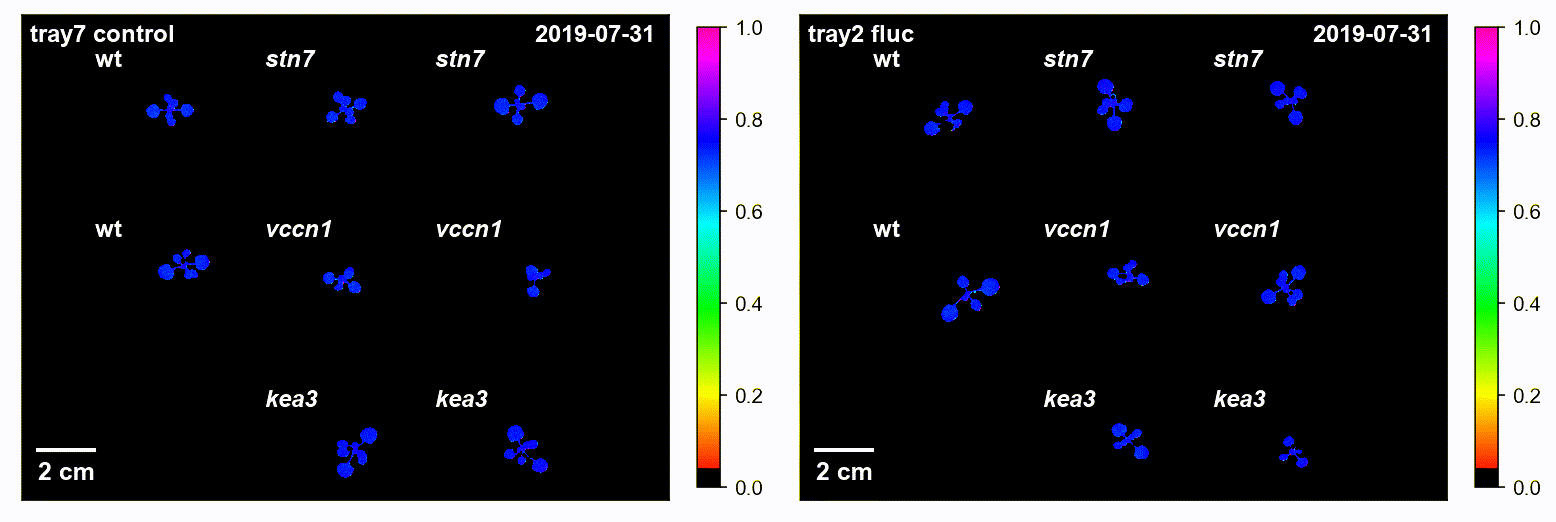

### FvFm_YII_tray7_x_tray3.gif

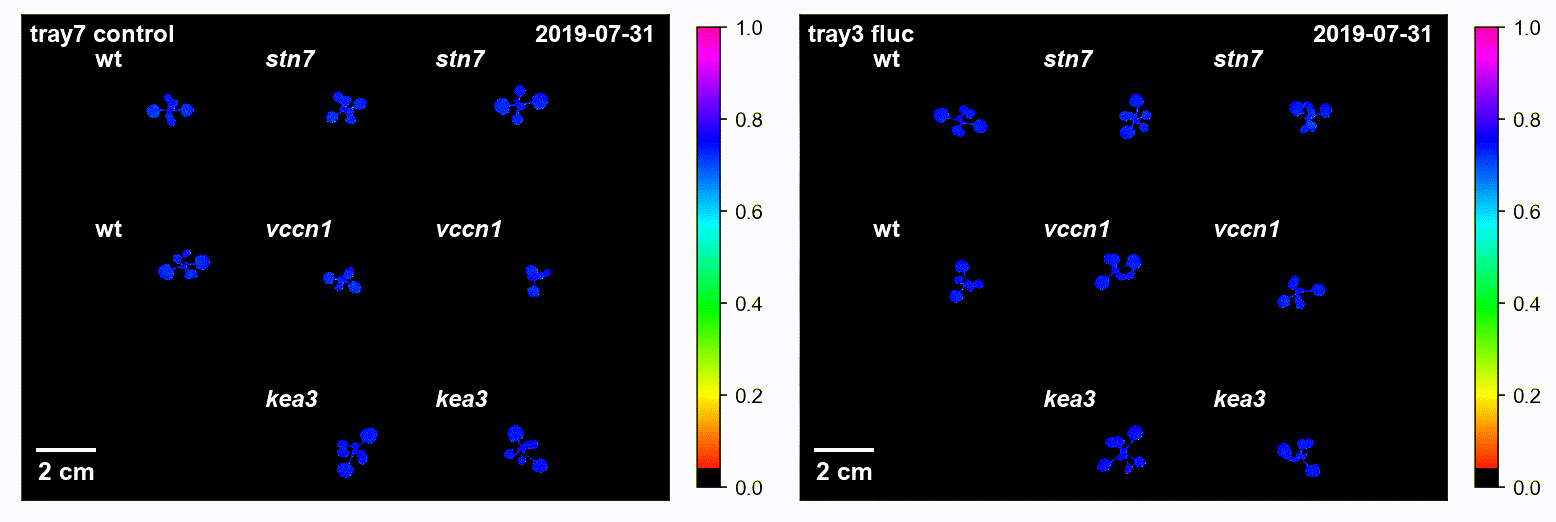

### FvFm_YII_tray7_x_tray4.gif

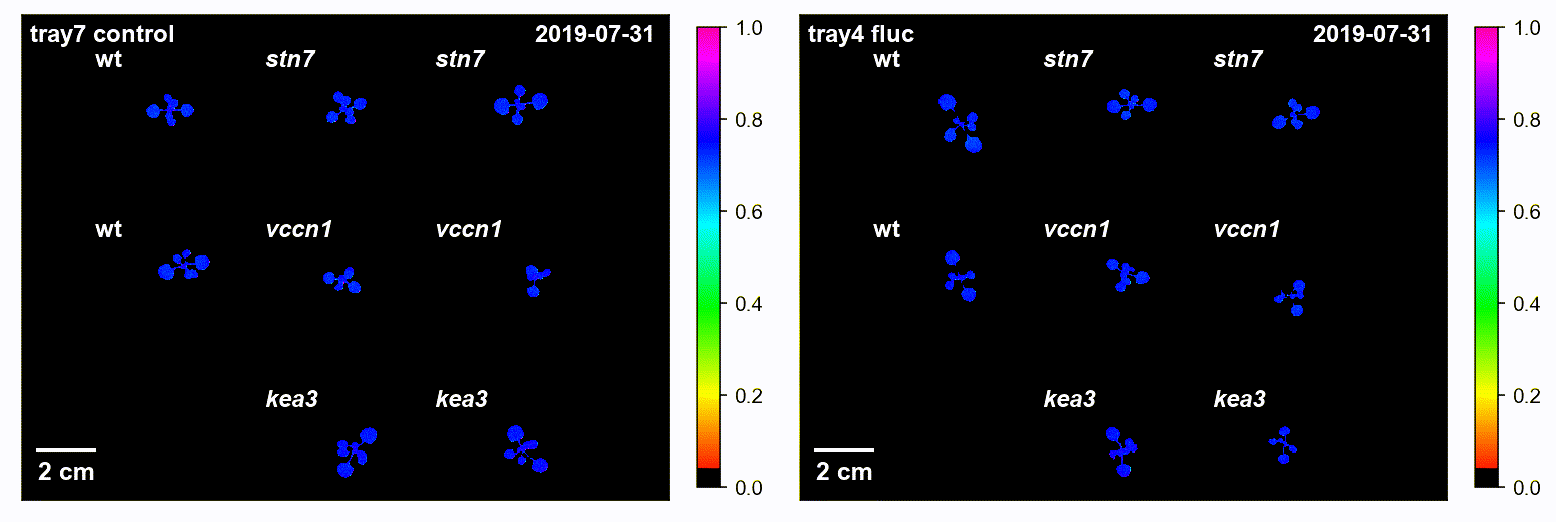

### t300_ALon_NPQ_tray5_x_tray2.gif

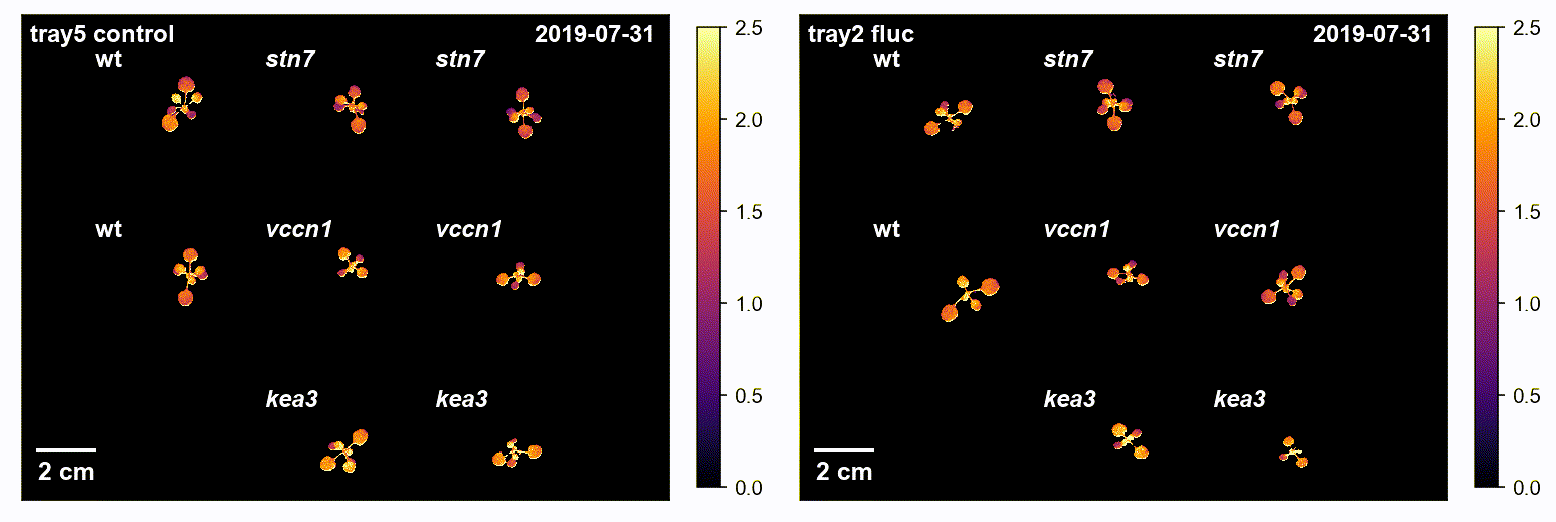

### t300_ALon_NPQ_tray5_x_tray3.gif

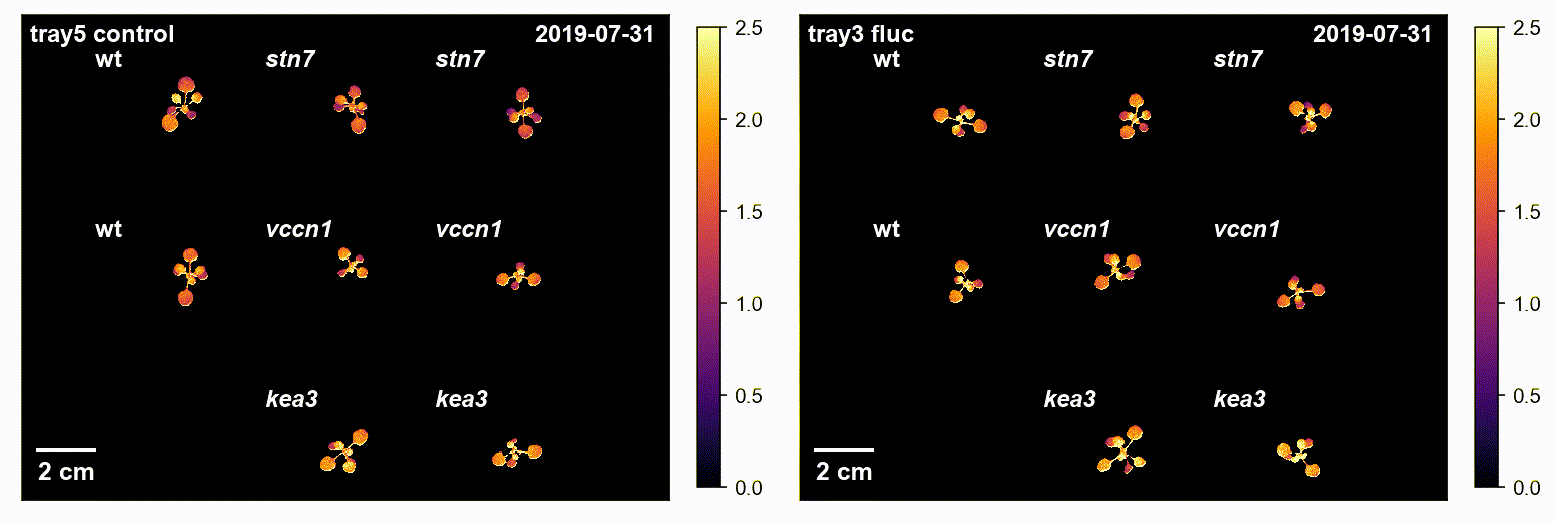

### t300_ALon_NPQ_tray5_x_tray4.gif

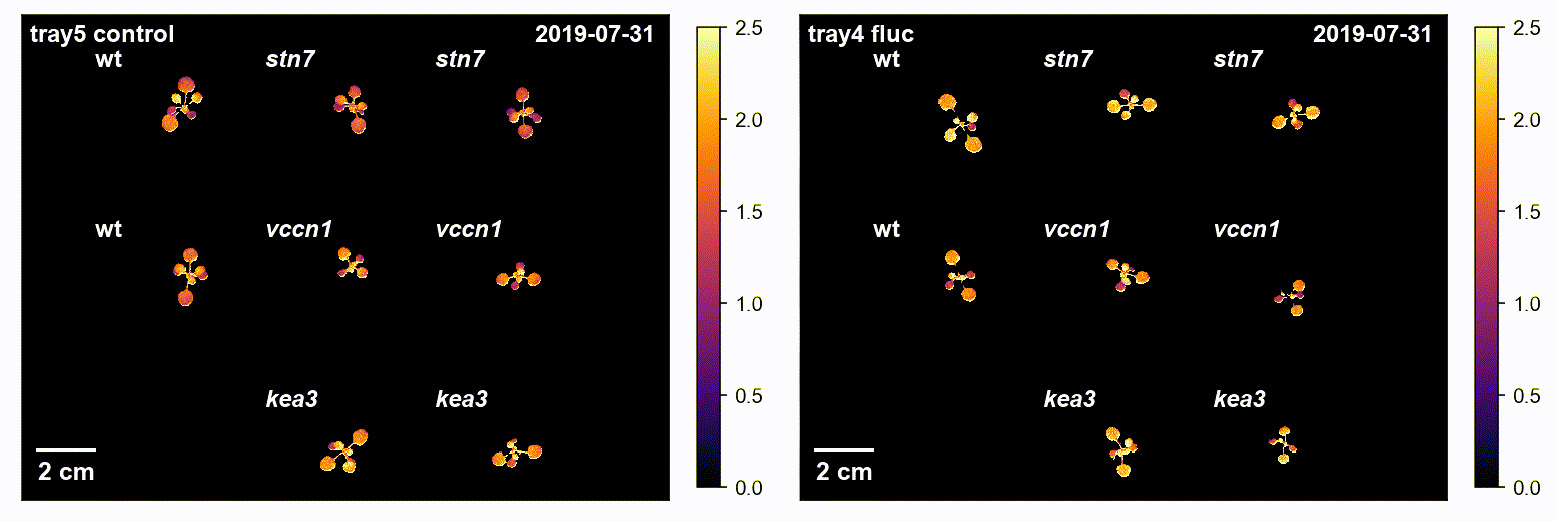

### t300_ALon_NPQ_tray6_x_tray2.gif

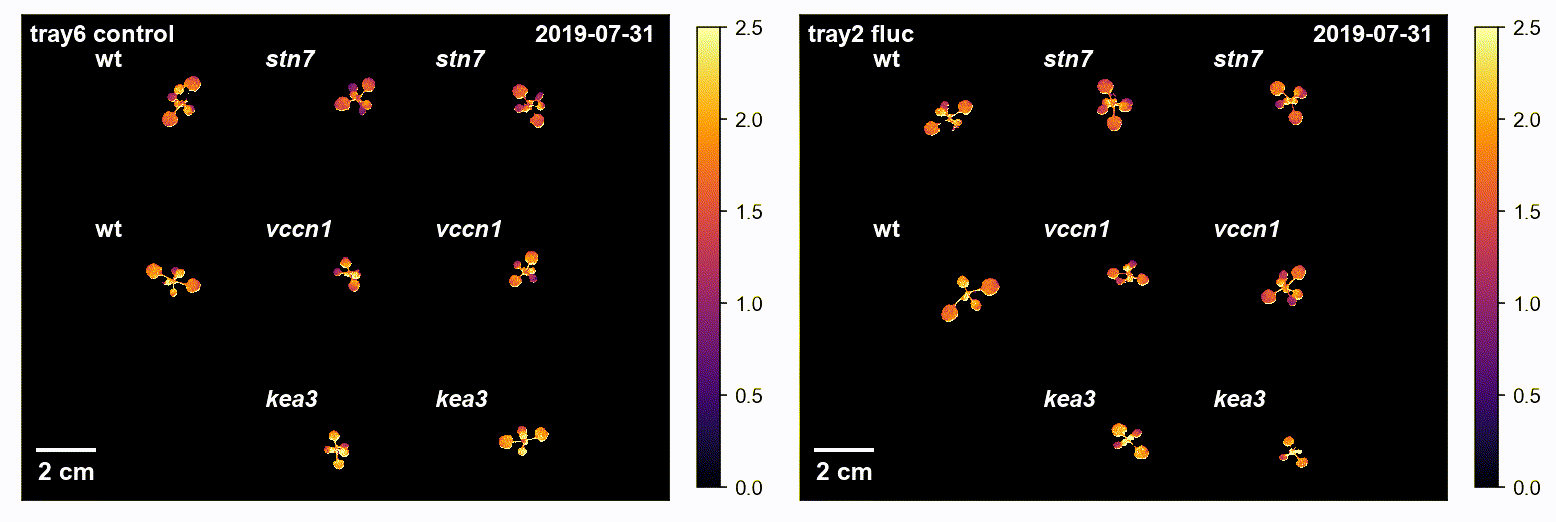

### t300_ALon_NPQ_tray6_x_tray3.gif

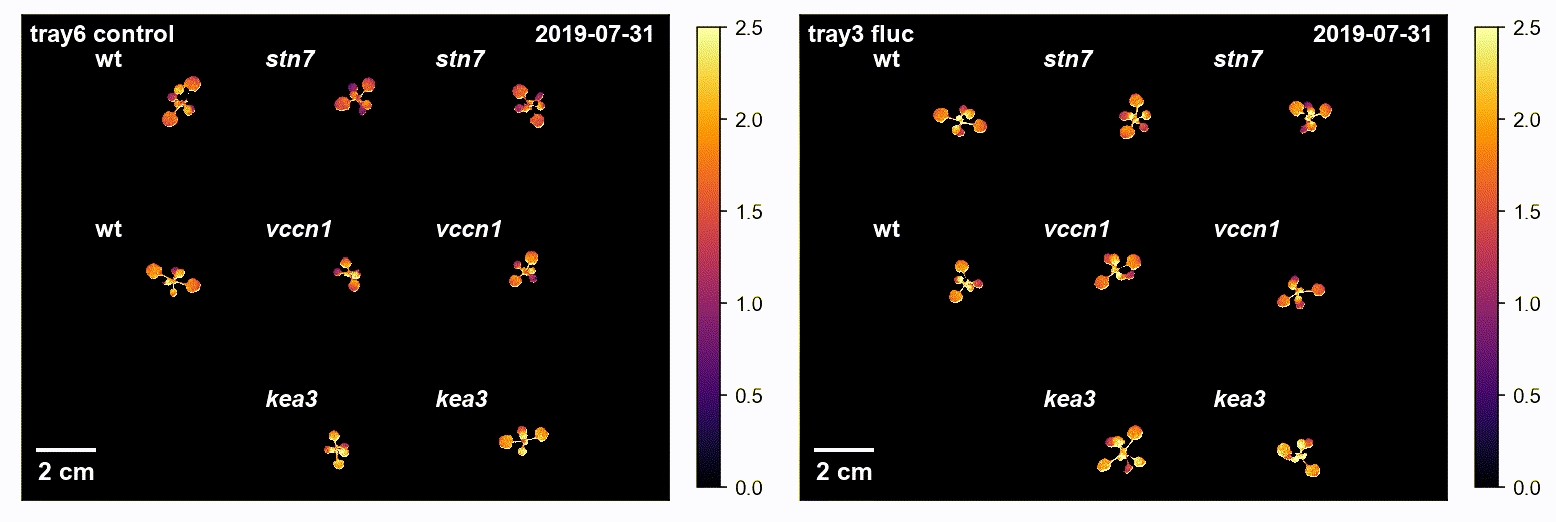

### t300_ALon_NPQ_tray6_x_tray4.gif

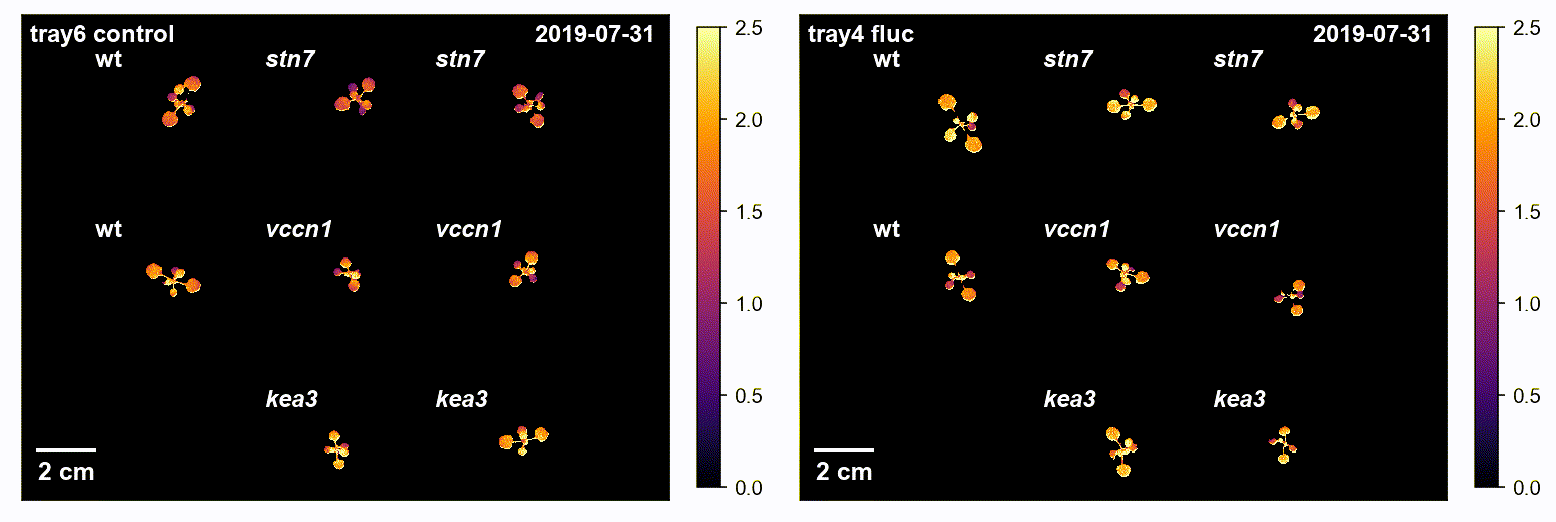

### t300_ALon_NPQ_tray7_x_tray2.gif

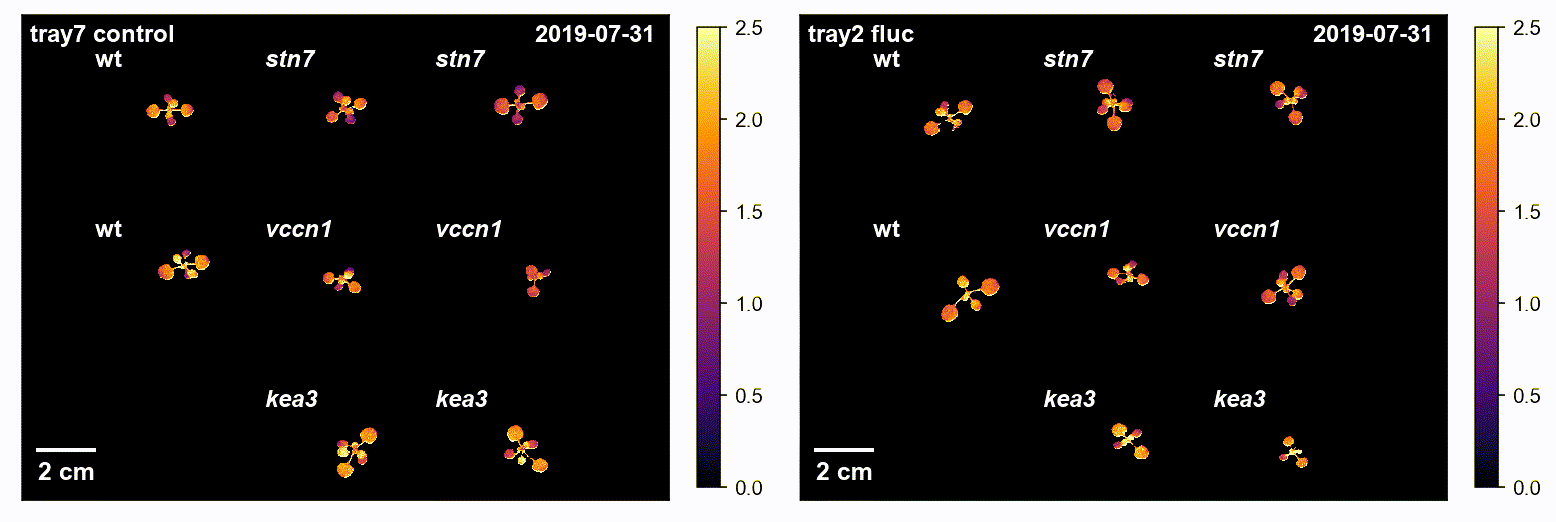

### t300_ALon_NPQ_tray7_x_tray3.gif

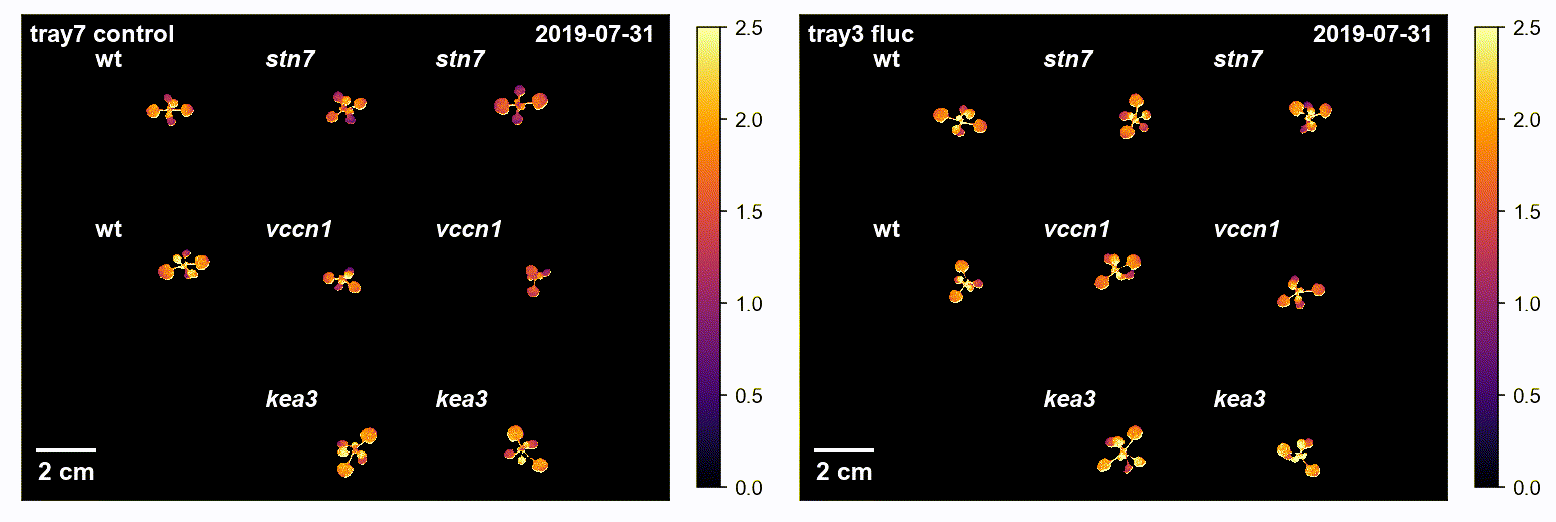

### t300_ALon_NPQ_tray7_x_tray4.gif

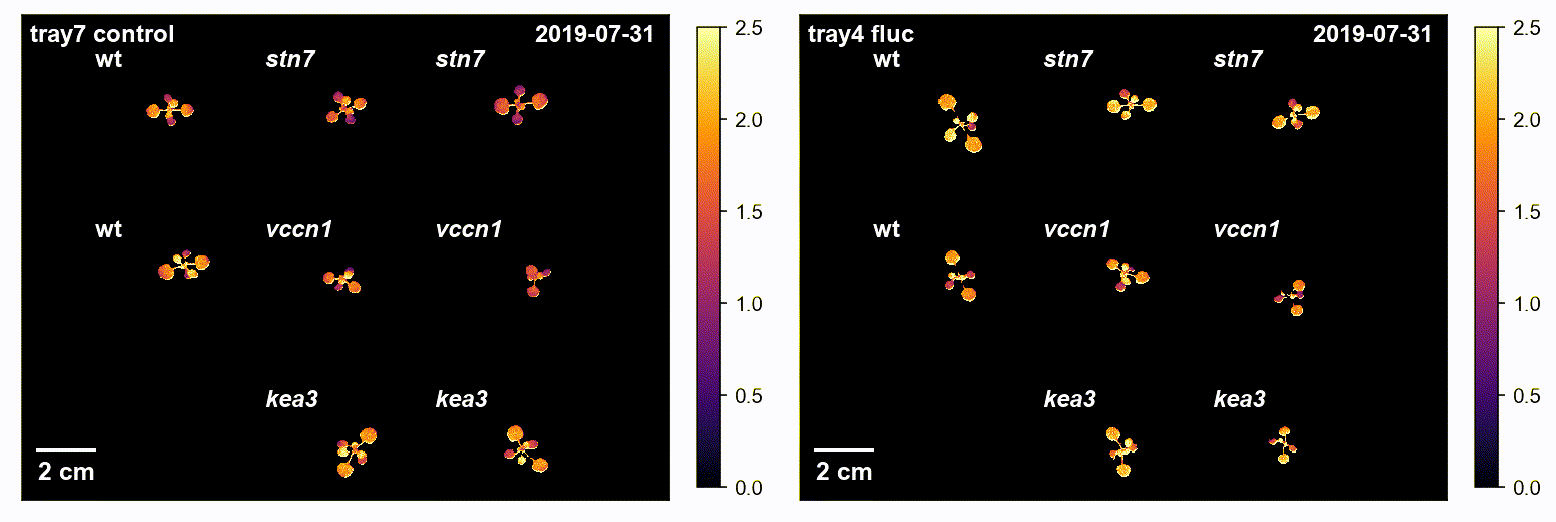

### t300_ALon_YII_tray5_x_tray2.gif

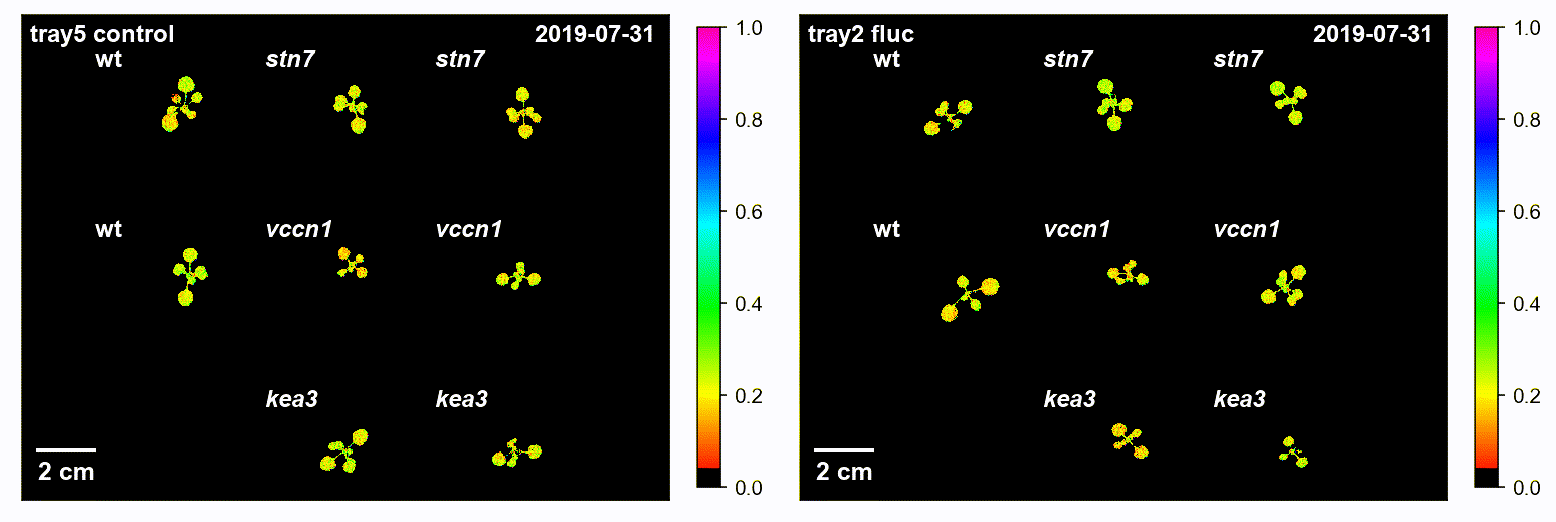

### t300_ALon_YII_tray5_x_tray3.gif

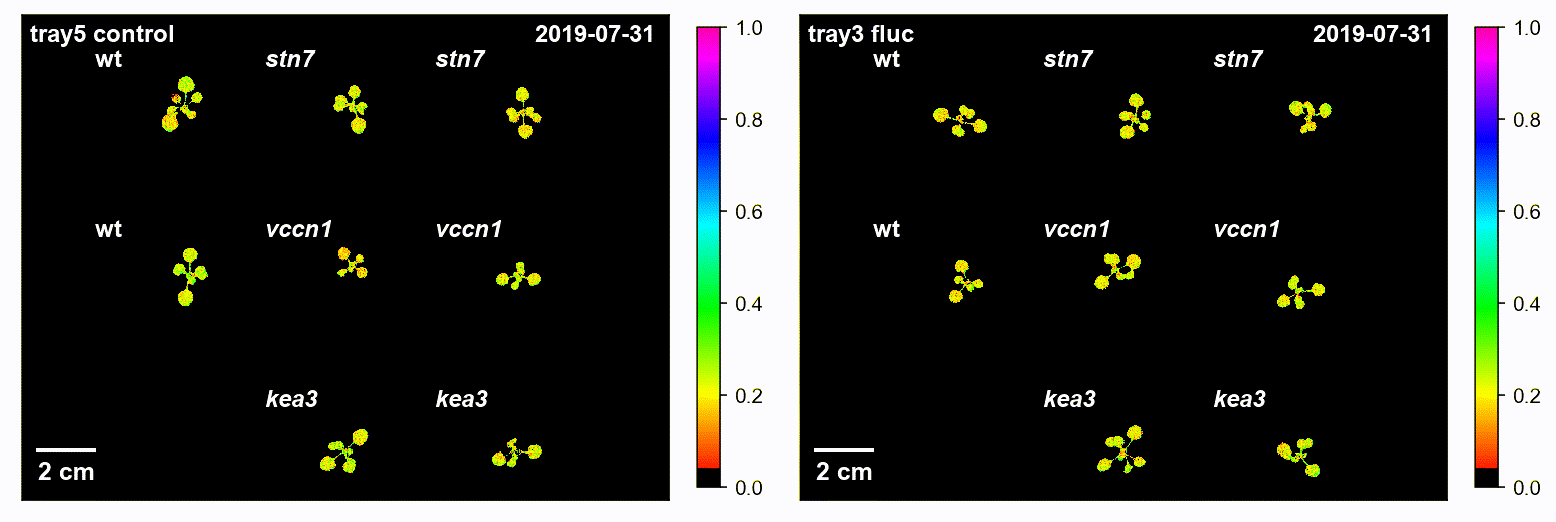

### t300_ALon_YII_tray5_x_tray4.gif

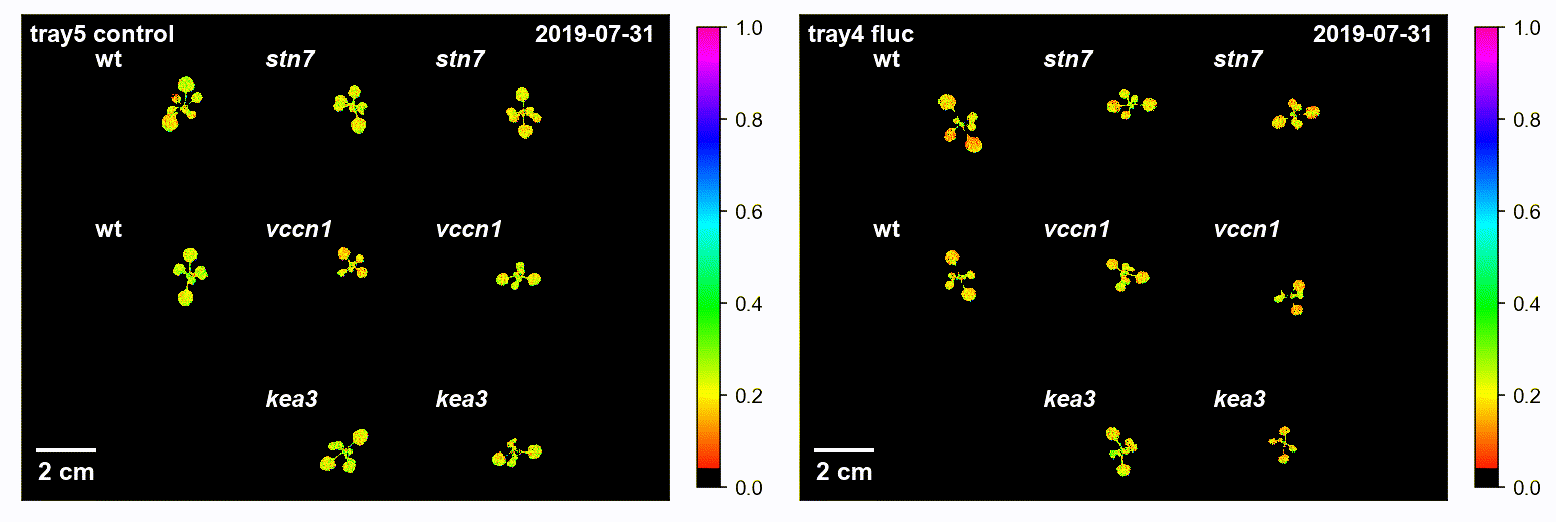

### t300_ALon_YII_tray6_x_tray2.gif

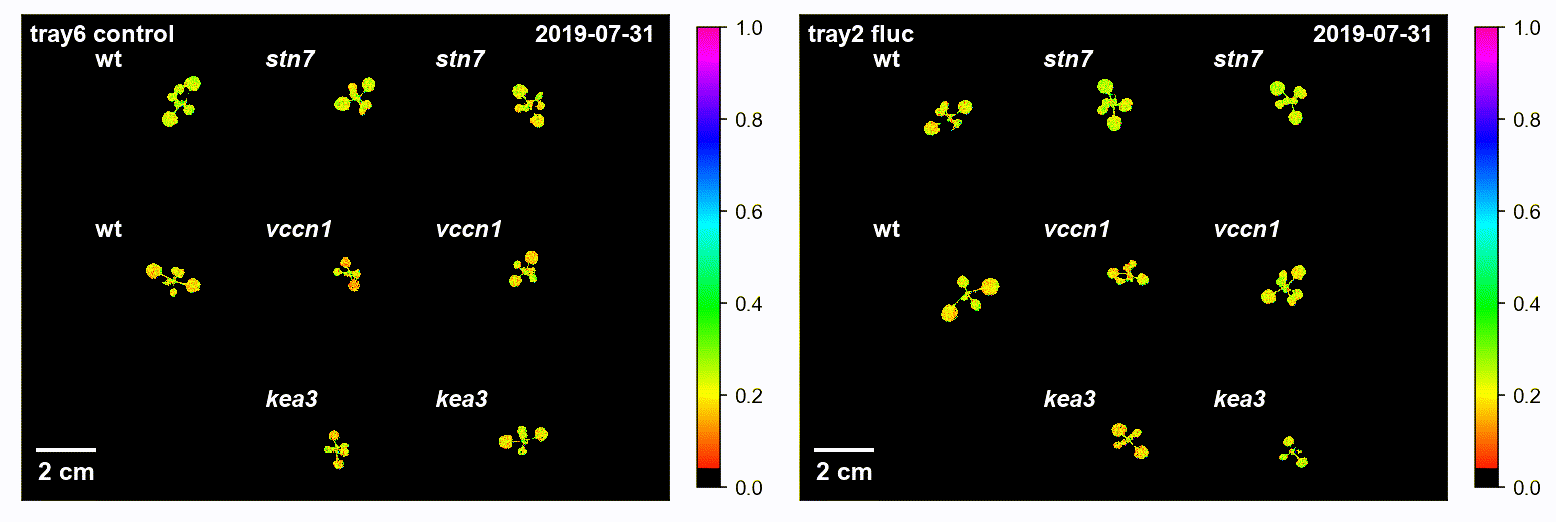

### t300_ALon_YII_tray6_x_tray3.gif

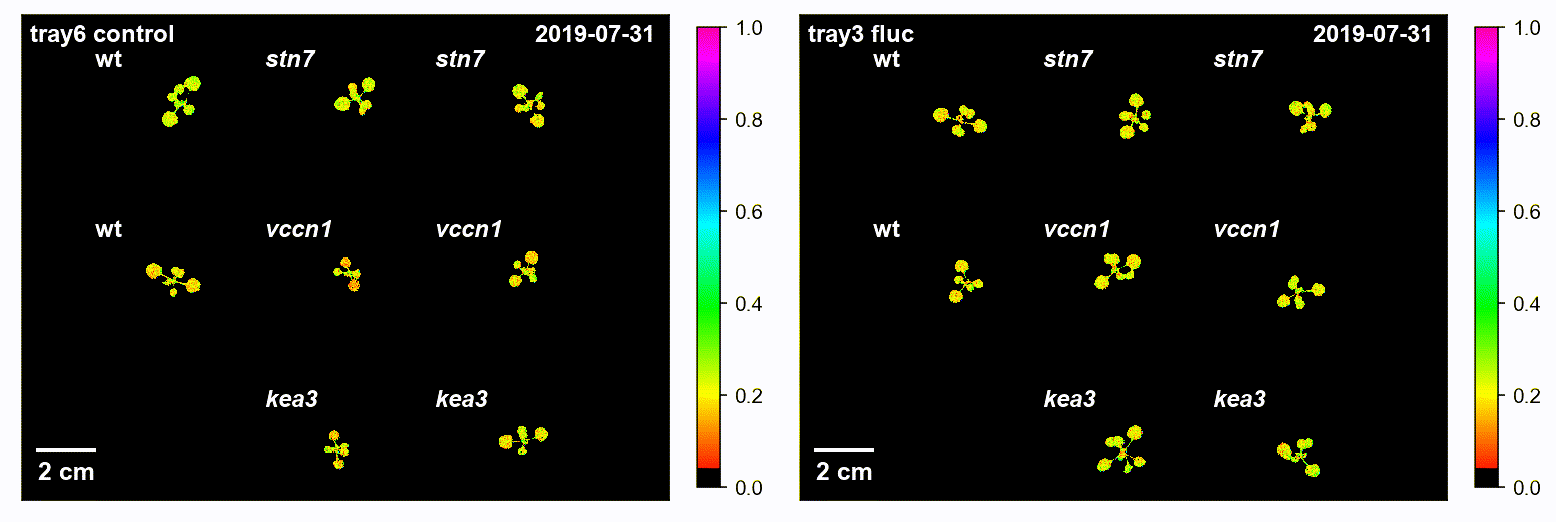

### t300_ALon_YII_tray6_x_tray4.gif

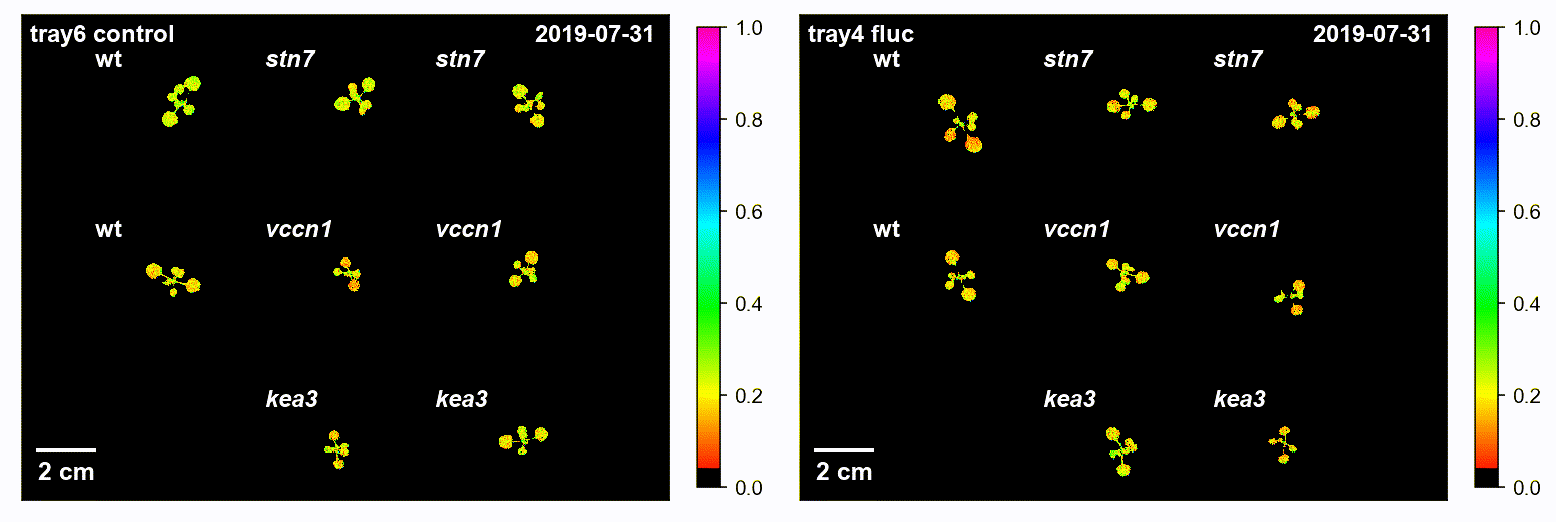

### t300_ALon_YII_tray7_x_tray2.gif

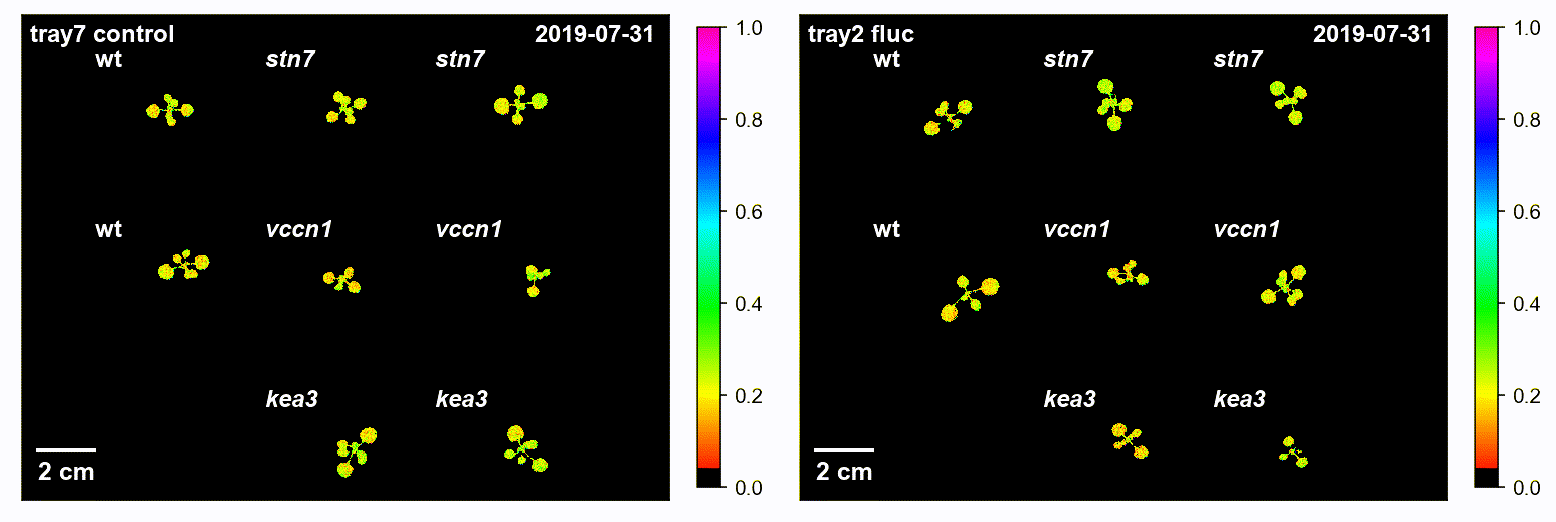

### t300_ALon_YII_tray7_x_tray3.gif

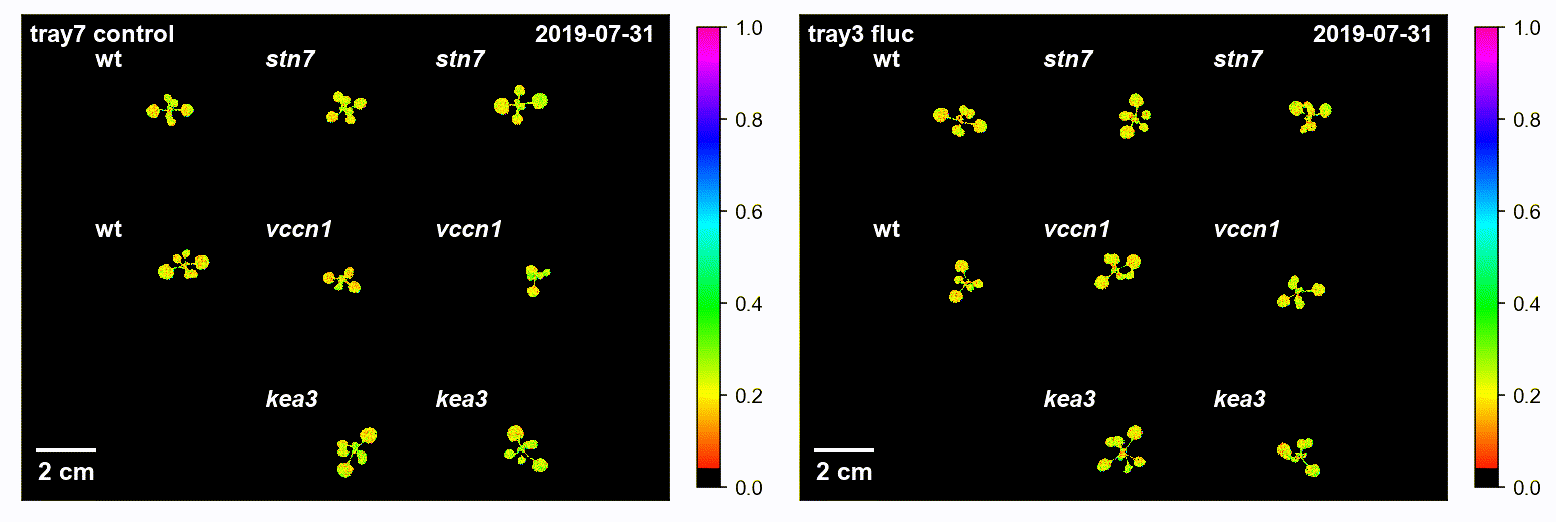

### t300_ALon_YII_tray7_x_tray4.gif

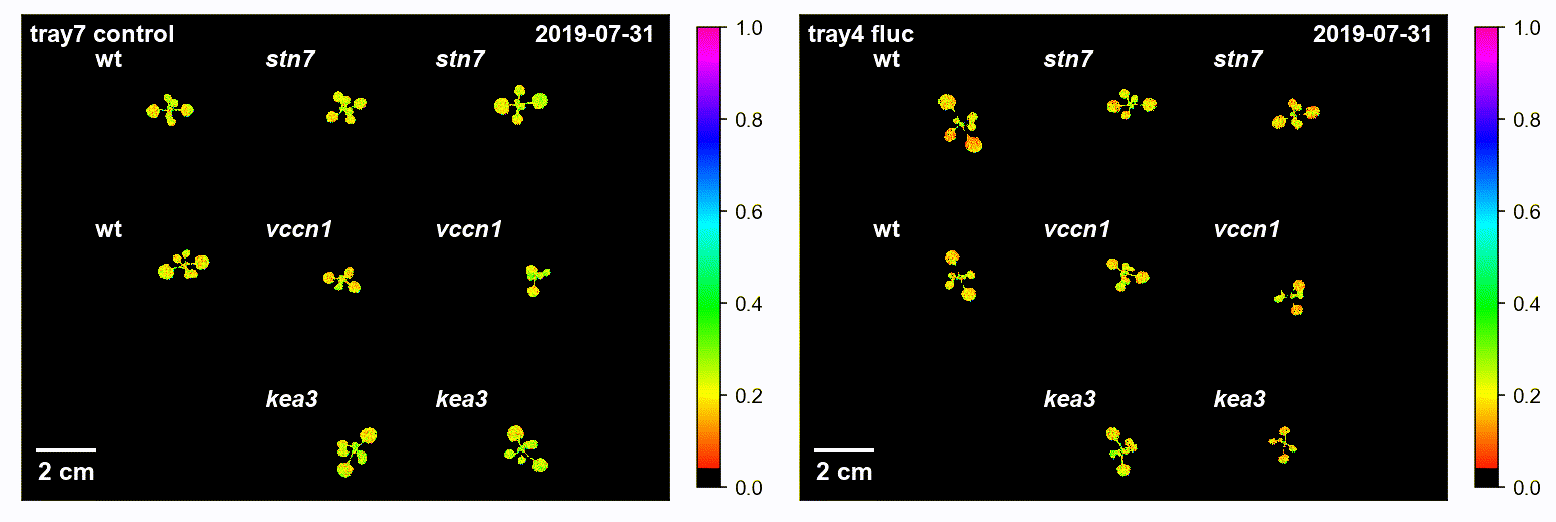
